# Lightning Pose 3D: an uncertainty-aware framework for data-efficient multi-view animal pose estimation

**DOI:** 10.64898/2026.04.20.719731

**Authors:** Lenny Aharon, Matthew R Whiteway, Karan Sikka, Keemin Lee, Yanchen Wang, Selmaan Chettih, Benjamin Midler, Ilana B Witten, Dmitriy Aronov, International Brain Laboratory, Liam Paninski

**Author notes:** These authors contributed equally to this work (Lenny Aharon, Matthew R Whiteway).

## Abstract

Multi-view pose estimation is essential for quantifying animal behavior in scientific research, yet current methods struggle to achieve accurate tracking with limited labeled data and suffer from poor uncertainty estimates. We address these challenges with a flexible framework that can operate with or without camera calibration, combining novel training and post-processing techniques with an uncertainty-aware pseudo-labeling distillation procedure. Our multi-view model processes all camera views jointly using a pretrained vision transformer backbone, and a simulated occlusion technique encourages the model to learn robust cross-view correspondences without requiring camera parameters. When camera parameters are available, 3D data augmentations and a triangulation-based loss further encourage geometric consistency. We extend the Ensemble Kalman Smoother (EKS) post-processor to the nonlinear case, leveraging camera geometry, and introduce a variance inflation technique that detects cross-view inconsistencies and corrects overconfident predictions. We validate our approach on five datasets spanning three species (fly, mouse, bird), including a multi-animal dataset with two visually distinct individuals; the proposed pipeline consistently outperforms existing methods across datasets. We demonstrate how these improvements translate to downstream scientific analyses using data from the International Brain Laboratory, showing improved unsupervised behavioral clustering and neural decoding of paw kinematics with just 200 labeled frames. To facilitate adoption, we developed a browser-based, cloud-compatible user interface that supports the full life cycle of multi-view pose estimation, from labeling and model training to post-processing with EKS and diagnostic visualizations.

## 1 Introduction

Precise measurement of animal behavior is central to modern neuroscience, and pose estimation—the automatic tracking of body part locations from video—has become a foundational tool for this purpose (Mathis et al., 2020; Pereira et al., 2020). A rich ecosystem of software packages now enables researchers to track user-defined keypoints with high accuracy in single-camera settings (Mathis et al., 2018; Graving et al., 2019; Pereira et al., 2019; Pereira et al., 2022; Biderman et al., 2024). However, single-camera approaches are fundamentally limited by occlusions and the inability to recover three-dimensional kinematics (Marshall et al., 2022; Moore et al., 2022). To address these limitations, a growing number of laboratories use multiple synchronized cameras to enable 3D pose reconstruction, which is critical for studying freely moving animals in complex arenas and during social interactions (Günel et al., 2019; Bala et al., 2020; Dunn et al., 2021; Karashchuk et al., 2021; Han et al., 2024; Cheng et al., 2025; Klibaite et al., 2025; Daruwalla et al., 2024).

The most common approach to multi-view pose estimation follows a modular pipeline: cameras are first calibrated (Fig. 1a,b), a view-agnostic 2D pose estimation network is trained on manually labeled frames (Fig. 1c,d), and predictions from each view are triangulated post hoc to obtain 3D coordinates (Fig. 1e) (Nath et al., 2019; Karashchuk et al., 2021; Maree et al., 2024). Although effective in certain settings, this approach has several important limitations. First, annotation cost scales linearly with the number of camera views, creating a strong need for methods that perform well with limited labeled data. Second, because each view is processed independently, the network must resolve occlusions and ambiguities in isolation, failing to exploit statistical regularities across views that could improve generalization to out-of-distribution data (Biderman et al., 2024). Third, precise camera calibration is difficult to obtain and maintain, and drift between sessions can bias 3D reconstructions. Fourth, network confidence scores can remain high even for incorrect predictions, providing no principled mechanism for identifying which frames require correction (Biderman et al., 2024).

**Figure 1:**
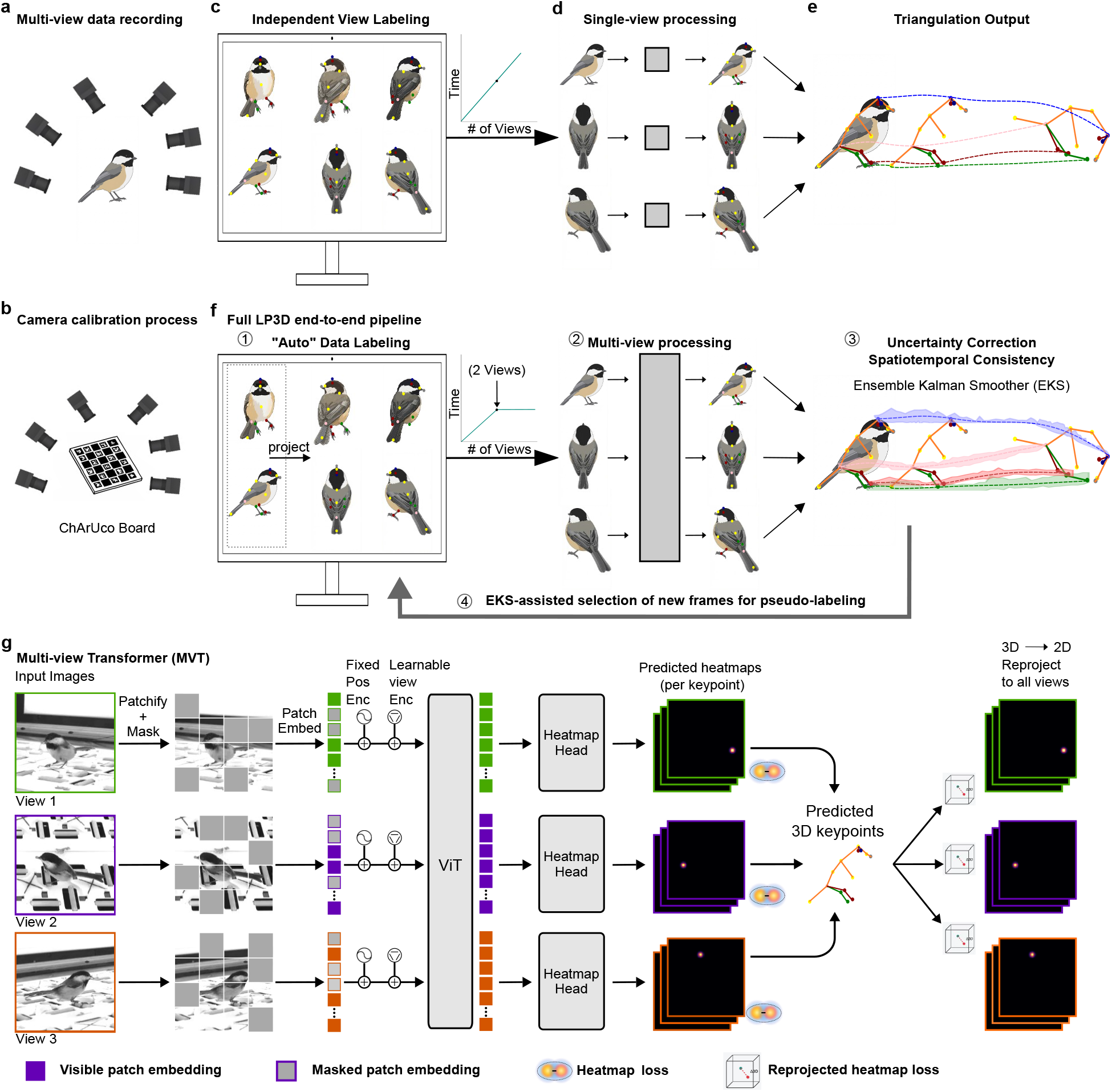
End-to-end pipeline for multi-view pose estimation. **a**, Data recording. **b**, The standard procedure for obtaining camera parameters involves recording a ChArUco board (a standard checkerboard-style calibration target) across all views (Karashchuk et al., 2021). **c**, Conventional multi-view pose estimation requires manual keypoint labeling in all views, with annotation cost scaling linearly with the number of views. **d**, Each view is independently processed with a view-agnostic 2D pose network. **e**, Per-view predictions are triangulated to obtain point estimates of 3D keypoints. **f**, Schematic of proposed framework. (1) Only two views are manually labeled; remaining views are automatically annotated via reprojection when calibration is available (Karaschuk, 2026). (2) A multi-view model processes all views jointly in a single forward pass, combining information across views directly during feature extraction, and operates with or without camera parameters. (3) A post-hoc uncertainty correction step produces improved predictions and uncertainty estimates, and operates with or without camera parameters. (4) Uncertainty estimates guide the selection of informative frames for pseudo-labeling, forming an iterative end-to-end training pipeline. **g**, Multi-view transformer (MVT) architecture. Pixel patches are randomly masked before patch embedding, then added to fixed positional and learnable view encodings. A single vision transformer (V_I_T) processes all views simultaneously. Training minimizes MSE loss between predicted and ground truth heatmaps. Multi-view consistency is additionally enforced via a 2D → 3D → 2D reprojection cycle: predicted keypoints are triangulated to 3D, reprojected back to 2D heatmaps, and compared against ground truth 2D heatmaps with an MSE loss.

Some methods move beyond this view-independent design; for example, DANNCE (Dunn et al., 2021) processes all views jointly in a learned volumetric representation. However, such methods require precise camera calibration as a prerequisite, so the calibration bottleneck often persists even for more sophisticated architectures. Multi-view human pose estimation has produced increasingly sophisticated architectures for this problem (Nogueira et al., 2025), but these methods are typically trained on large standardized datasets with fixed keypoint definitions. These resources do not exist for animal pose estimation, where labeled datasets are typically small (though see Marshall et al., 2021; Ulutas et al., 2025) and keypoint sets vary across species and experiments. Approaches designed for the animal setting must therefore achieve strong performance with limited labeled data while remaining flexible across diverse recording configurations.

In addition to methodological limitations, the software ecosystem for multi-view pose estimation is fragmented, with no single solution covering all stages from calibration through network training to post-processing. This means each laboratory must assemble and debug its own pipeline by stitching together separate tools, requiring careful engineering. Iterating on performance—identifying pose estimation errors, labeling new data, retraining networks, and rerunning inference—is similarly labor- and skill-intensive. The practical consequence is that researchers may stop iterating before pose estimation quality is sufficient, allowing residual errors to negatively affect downstream analyses.

To address these shortcomings we introduce Lightning Pose 3D (LP3D), an end-to-end framework for multi-view animal pose estimation that exploits cross-view geometric consistency at every stage of the pipeline (Fig. 1f). During training, a multi-view transformer jointly processes all views, and a novel simulated occlusion approach learns cross-view correspondences without requiring camera calibration. When calibration is available, 3D data augmentation and a triangulation-based loss further encourage geometric consistency.

For post-processing, we extend the Ensemble Kalman Smoother (EKS) introduced in Biderman et al. (2024) to the nonlinear setting, enabling it to leverage camera geometry when available. A novel variance inflation technique for EKS detects cross-view inconsistencies and corrects overconfident predictions. The resulting calibrated uncertainties drive a pseudo-labeling pipeline that automatically selects high quality labels for retraining, enabling iterative improvement with minimal additional annotation.

We validate our approach on five datasets spanning three different species (flies, mice and chickadees), including a multi-animal dataset with two visually distinct mice. We show that each component of the framework contributes complementary benefits, and that the full pipeline consistently outperforms existing methods across datasets. We further demonstrate that these improvements in pose accuracy lead to cleaner behavioral representations and improved decoding of behavioral variables from neural activity in a dataset from the International Brain Laboratory (IBL et al., 2025). We release an open-source software package and browser-based, cloud-compatible user interface that supports the full life cycle of multi-view pose estimation, from data labeling and model training to post-processing with EKS and diagnostic visualization.

## 2 Results

### 2.1 A flexible and data-efficient multi-view transformer (MVT) pose estimation model

Most multi-view pose estimation pipelines process each camera view independently and fuse predictions post hoc via triangulation, requiring each view to resolve occlusions and ambiguities in isolation. Our multi-view transformer architecture (MVT) addresses this fundamental limitation by combining information from all views early in processing: image patches from all camera views are combined into a single input and processed together by a pretrained computer vision backbone in one pass (Fig. 1g). Each patch is tagged with its spatial location within the image and which camera it came from, without requiring explicit camera parameters. This allows the model to make predictions about a body part that is occluded in one view by using information about its location from non-occluded views.

We designed the MVT data processing to be compatible with a range of pretained vision transformer back-bones. This flexibility allows us to update the backbone as foundation models improve, and also allows us to select large backbones when accuracy is more important than speed, and small backbones when speed is more important than accuracy. For our analysis, we use a small (∼ 21M parameters) vision transformer (ViT) pretrained with the DINO self-supervised learning objective (Caron et al., 2021) as the default backbone, balancing accuracy and speed (Supp Figs. 1, 2).

These design choices translate to strong performance: our approach outperforms independent per-view networks, including DeepLabCut (Mathis et al., 2018), across all datasets (Extended Data Fig. 1a, 2). We apply the same training procedure and hyperparameters across datasets, demonstrating a robustness that is important for adoption in experimental labs where extensive hyperparameter tuning is impractical (Table 1).

**Table 1:** Training hyperparameters by dataset. All datasets share the same architecture (multi-view transformer with CNN head), optimizer (Adam), and most training settings. Dataset-specific differences are highlighted where applicable. Entries correspond to parameter fields in the standard Lighting Pose model configuration file (see example at https://github.com/paninski-lab/lightning-pose/blob/v2.0.8/scripts/configs/config_default_multiview.yaml).

| Hyperparameter | Dataset |  |  |  |  |
| --- | --- | --- | --- | --- | --- |
|  | Treadmill Mouse | Fly | Chickadee | IBL | Resident-Intruder |
| config.data |  |  |  |  |  |
| image_resize_dims |  |  | 256×256 |  |  |
| num_keypoints | 7 | 30 | 18 | 3 | 22 |
| config.model |  |  |  |  |  |
| backbone |  |  | vits_dino |  |  |
| model_type |  |  | heatmap_multiview_transformer |  |  |
| head |  |  | heatmap_cnn |  |  |
| heatmap_loss_type |  |  | mse |  |  |
| config.training |  |  |  |  |  |
| optimizer |  |  | Adam |  |  |
| optimizer_params.learning_rate |  |  | 5e-5 |  |  |
| lr_scheduler_params.multisteplr.gamma |  |  | 0.5 |  |  |
| lr_scheduler_params.multisteplr.milestones |  |  | [2000, 3000, 4000] |  |  |
| train_batch_size |  |  | 8 |  |  |
| train_prob / val_prob |  |  | 0.95 / 0.05 |  |  |
| imgaug |  |  | dlc |  |  |
| imgaug_3d | false | true | true | true | true |
| patch_mask.init_step |  |  | 700 |  |  |
| patch_mask.final_step |  |  | 5000 |  |  |
| patch_mask.init_ratio |  |  | 0.1 |  |  |
| patch_mask.final_ratio |  |  | 0.5 |  |  |
| unfreezing_step |  |  | 400 |  |  |
| max_steps |  |  | 5000 |  |  |
| config.losses |  |  |  |  |  |
| supervised_reprojection_heatmap_mse.log_weight | — | 3 | 3 | 3 | 3 |
| config.callbacks |  |  |  |  |  |
| anneal_weight.freeze_until_epoch | — | 60 | 60 | 60 | 60 |

### 2.2 Robust MVT training with simulated occlusions and geometric consistency loss (LP3D)

To further improve robustness, we address two remaining failure modes. First, although joint multi-view processing reduces the ambiguity of single-view methods, occlusions are not fully eliminated. When training on a limited set of labeled frames, the model is exposed to only a small fraction of the occlusion patterns that can arise in practice, leaving it vulnerable to novel occlusions at test time.

To address this, we introduce patch masking: during training, contiguous patches are randomly masked in pixel space, simulating the large occlusions that arise in videos (Fig. 1g; Supp Fig. 3). Because the masked keypoints often cannot be resolved from the affected view alone, the model can route attention towards other views where the body parts remain visible. Since masking patterns are sampled randomly, each labeled instance presents a distinct configuration of visible and occluded regions each time it is seen during training, forcing the model to learn flexible cross-view attention rather than memorizing fixed occlusion patterns. Patch masking requires no camera parameters and can therefore be applied to any dataset.

A second, independent failure mode concerns geometric consistency. Without explicit geometric constraints, per-view 2D predictions can be individually plausible, but a body part triangulated from different view pairs may resolve to different 3D locations. This means the network can appear to perform well on each view in isolation while producing predictions that are physically impossible when considered jointly.

When camera parameters are available, we address this with geometrically consistent 3D data augmentation across all views (Supp Figs. 4, 5) and a 3D reprojection loss: predicted keypoints are triangulated to 3D, reprojected back to 2D heatmaps, and compared against the ground truth 2D heatmaps (Fig. 1g). A single hyperparameter weights this loss against the 2D heatmap loss, and is set to the same value across all calibrated datasets, without per-dataset tuning (Supp Fig. 6). For uncalibrated setups (e.g., Treadmill Mouse), the 3D augmentations and reprojection loss are simply omitted without any change to the training procedure.

Patch masking and the reprojection loss address distinct failure modes and are therefore complementary. Each individually outperforms the base MVT, and their combination achieves the best performance across all datasets (Supp Fig. 7). We define the MVT with patch masking and the reprojection loss as LP3D.

### 2.3 LP3D improves pose estimation performance

To assess whether LP3D generalizes across species, recording configurations, and calibration conditions, we evaluated it on five datasets spanning three species (Fig. 2a,b): a head-fixed mouse on a treadmill recorded with a mirrored setup, resulting in two views (“Treadmill Mouse”; Warren et al., 2021); a head-fixed fly recorded from six cameras (“Fly”; Karashchuk et al., 2021); a freely moving chickadee recorded from six cameras (“Chickadee”; Chettih et al., 2024); a head-fixed mouse performing a decision-making task recorded from two cameras, from the International Brain Laboratory (“IBL”; IBL et al., 2025); and two interacting, visually distinct mice recorded from five cameras (“Resident-Intruder”).

**Figure 2:**
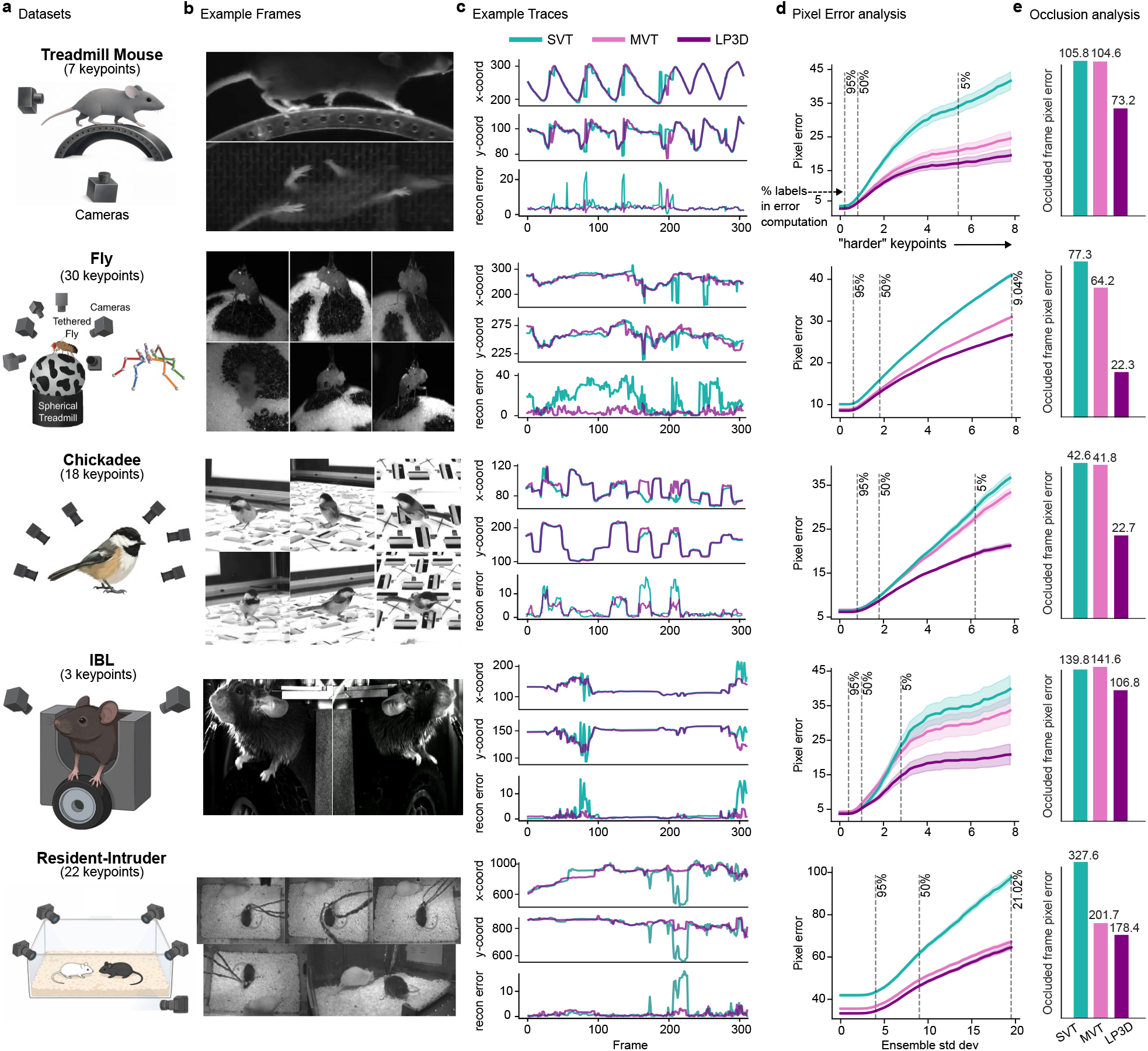
LP3D improves pose estimation across diverse datasets. **a**, Experimental setups and labeled keypoints for all five datasets. **b**, Example frames from a single instance for each dataset. **c**, Example keypoint coordinate traces (top) and 3D reprojection error (bottom) for the single-view transformer (SVT; teal) and LP3D (purple). 3D reprojection error utilizes camera parameters to measure the consistency of predictions across camera views, requiring no ground truth labels. For Treadmill Mouse, which lacks camera parameters, consistency is measured via 3D PCA. LP3D produces more temporally consistent predictions and lower reprojection error across all datasets. **d**, Pixel error as a function of keypoint difficulty (lower is better), computed on frames from held-out test animals (Biderman et al., 2024). We computed the standard deviation of each keypoint prediction in each frame in the test data across all model types and seeds. We then took the mean pixel error over all keypoints with a standard deviation larger than a threshold value, for each model type. Dashed vertical lines indicate the percentage of data used for the pixel error computation. Error bands represent the s.e.m. over all included keypoints and frames for a given standard deviation threshold. SVT (teal), MVT without patch masking or reprojection loss (MVT; pink), and LP3D (purple) are shown for all datasets trained with 200 labeled frames. **e**, Held-out view analysis. At test time, one camera view is fully masked and models must predict keypoints in that view using the remaining views. SVT error represents chance since this model does not have access to additional information. Pixel error on the held-out view is shown for SVT, MVT, and LP3D, demonstrating that patch masking and the reprojection loss trains the model to leverage complementary information across views and better handle occlusions.

We train models with 200 labeled frames unless otherwise noted, and evaluate on frames from held-out test animals (Supp Videos 1-6). Our primary baseline is a single-view transformer (SVT) that shares the same ViT backbone as MVT but processes each view independently (Fig. 1d; Supp Figs. 8); the ViT backbone alone already yields improvements over prior ResNet-based approaches (Supp Fig. 1). Since SVT and MVT differ only in whether views are processed jointly, any performance gap directly measures the benefit of cross-view attention. Similarly, any performance gap between MVT and LP3D measures the contribution of patch masking and the reprojection loss.

LP3D consistently outperforms both SVT and MVT across all datasets. Keypoint traces show that LP3D produces more temporally consistent predictions than SVT, and 3D reprojection error is substantially lower for LP3D (Fig. 2c), where reprojection error measures how well predictions across views agree on a single 3D location (using 3D PCA for Treadmill Mouse, which lacks camera parameters). Quantitatively, LP3D achieves the lowest pixel error across all datasets and keypoint difficulty levels (Fig. 2d; Methods).

To directly test whether LP3D has learned genuine cross-view consistency rather than per-view shortcuts, we conducted a held-out view experiment: at test time, one camera view is fully masked and the model must predict keypoints in that view using only the remaining views. LP3D substantially outperforms both SVT and MVT in this setting (Fig. 2e), demonstrating that multi-view processing alone is not enough to leverage the power of cross-view attention in the MVT; the additional training objectives are required to drive the model to encode shared structure across views rather than relying on independent per-view features.

To test generalization across different data regimes, we trained models on subsets of 100 and 400 labeled frames. SVT generally outperforms or matches a ResNet-50, and LP3D maintains its advantage over all baselines even with only 100 labeled frames. Notably, the performance gap between LP3D and SVT/ResNet-50 models generally increases with more training data (Extended Data Fig. 2). Beyond label efficiency, LP3D’s accuracy scales with the number of training views, suggesting that richer multi-view supervision leads to stronger pose representations (Supp Fig. 9). LP3D also remains robust to the camera position shifts that naturally arise across recording sessions in experimental settings, maintaining strong performance despite session-to-session variability in camera placement and angle (Supp Fig. 10).

We also compared LP3D to DANNCE (Dunn et al., 2021), which constructs a learned volumetric representation from multi-view inputs. We first trained an LP3D model on a large-scale rat motion capture dataset (Rat7M; Marshall et al., 2021) using just 455 labeled frames—a number representative of what an individual lab might reasonably collect—and compared it to a DANNCE model trained on over 30,000 frames from the same dataset. While DANNCE outperforms LP3D in this setting, LP3D achieves strong performance with roughly 65× fewer labels (Extended Data Fig. 1b). To compare the two approaches under matched data conditions, we fine-tuned a DANNCE model pretrained on Rat7M on the Fly and Chickadee datasets using the same 200 labeled frames as LP3D. On both datasets, LP3D shows a substantial reduction in pixel error across all keypoint difficulty levels (Extended Data Fig. 1c; Supp Videos 7, 8).

### 2.4 A nonlinear post-processor with cross-view inconsistency detection

LP3D produces strong per-frame 2D predictions, but two limitations remain. First, individual predictions still contain jitter or errors on challenging frames (e.g., occlusions and motion blur). Second, although the reprojection loss encourages geometric consistency during training, it does not enforce it at inference/test time: predictions across views are not guaranteed to be explained by a single 3D point, and naively triangulating inconsistent predictions can amplify rather than correct errors. A post-processing step that jointly exploits temporal continuity and multi-view geometry, while providing calibrated uncertainty estimates that flag unreliable predictions, can address both limitations without retraining.

The multi-view Ensemble Kalman Smoother (mvEKS) introduced in Biderman et al., 2024 provides exactly this framework. Keypoints are modeled as noisy projections of a 3D latent state that evolves smoothly over time; an ensemble of networks provides both the input observations (median over networks) and initial uncertainty estimates (variance over networks). The smoother leverages temporal continuity and geometric consistency to produce corrected predictions with posterior uncertainty estimates (Fig. 3a). In Biderman et al., 2024, multi-view observations were modeled in a learned linear latent subspace that approximates shared 3D structure without explicit camera geometry (Fig. 3b; Extended Data Fig. 3c). Here we extend mvEKS to the nonlinear setting: when camera parameters are available, we use them to directly project the 3D latent to each view, enforcing the true geometric relationship rather than a linear approximation. The nonlinearversion outperforms the linear version on calibrated datasets (Extended Data Fig. 3d). For uncalibrated setups (e.g., Treadmill Mouse), the linear version is used. The only free hyperparameter is the smoothing strength, and we introduce an automatic algorithm to select this value per keypoint, avoiding manual tuning and recovering the correct temporal structure across datasets (Fig. 3c; Extended Data Fig. 3a,b).

**Figure 3:**
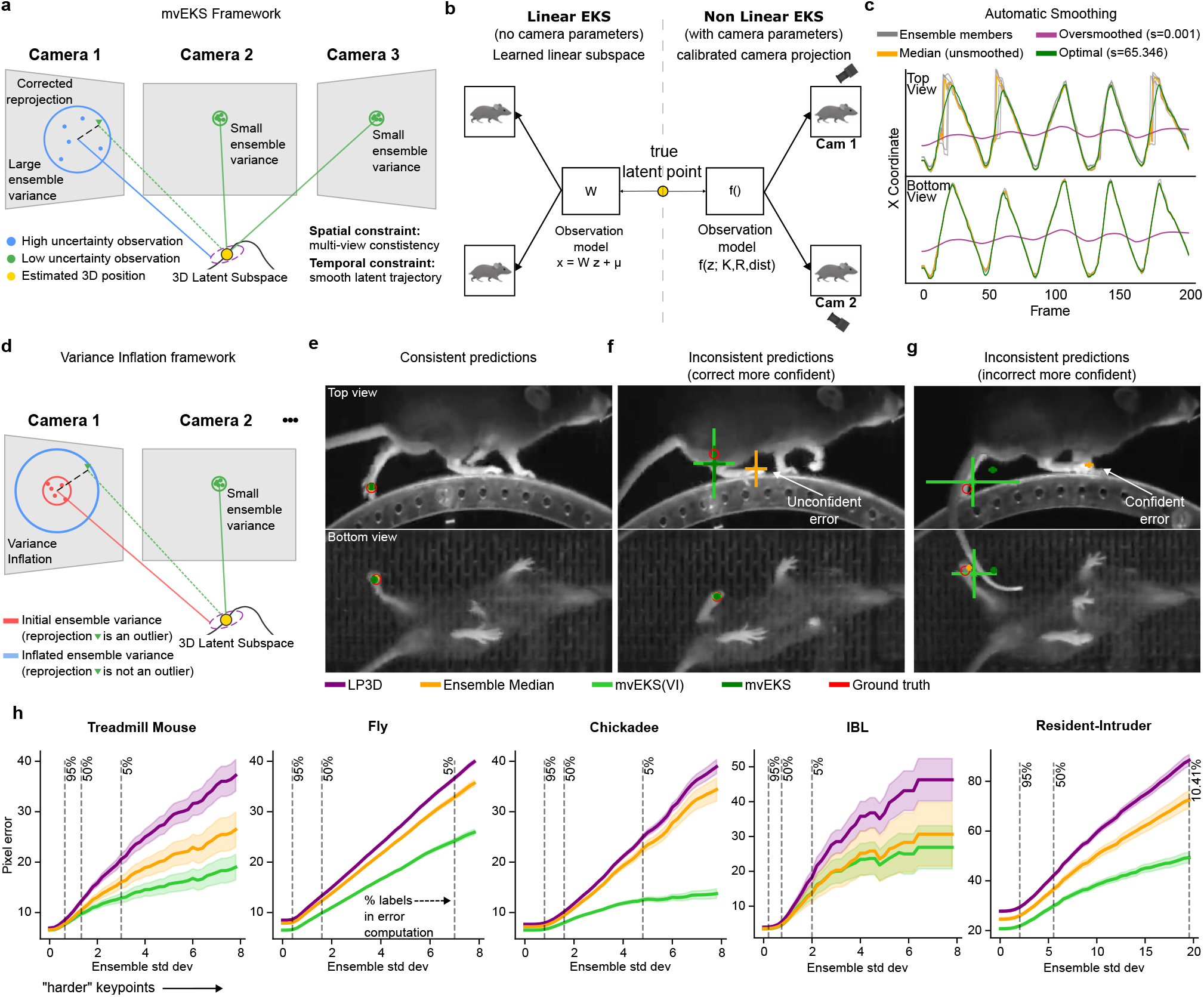
Multi-view Ensemble Kalman Smoother (mvEKS) improves pose estimation. **a**, EKS models keypoints as projections from a 3D latent that evolves smoothly over time; low-uncertainty observations from reliable camera views help correct high-uncertainty observations through spatial and temporal constraints. **b**, Linear vs. nonlinear EKS. *Left:* Without camera parameters, observations are modeled in a learned linear latent subspace. *Right:* With camera parameters, a nonlinear projection maps the 3D latent to each view, enforcing geometric consistency. **c**, Treadmill Mouse paw traces from two views. Optimal smoothing (green) recovers true oscillatory motion in the partly occluded Top view; oversmoothing (purple) distorts temporal dynamics. **d**, Variance inflation. For a given view and keypoint, if the reprojection from the remaining views is an outlier in the context of the ensemble variance, the ensemble is likely overconfident in that view; the variance is increased until the reprojection is no longer an outlier, down-weighting its influence on the EKS posterior. **e**, Consistent predictions across views; no variance inflation required. **f**, Inconsistent predictions across views detected by variance inflation, where the more confident predictions are correct. Orange crosses are ensemble median with ensemble variance; green crosses are corrected predictions from mvEKS with posterior predictive variance. **g**, Inconsistent predictions between views where a highly confident but incorrect prediction in the top view dominates; mvEKS is unable to override the confident error, but the variance inflation procedure adjusts the posterior predictive variance to reflect the remaining uncertainty. **h**, Ensemble median (orange) outperforms individual LP3D models (purple); nonlinear variance-inflated mvEKS (light green) achieves the best performance. Treadmill mouse (uncalibrated setup) uses linear mvEKS.

A subtler failure mode arises when ensemble members agree on an incorrect prediction. Standard EKS assumes that ensemble variance faithfully reflects prediction reliability. But when all ensemble members are confidently wrong, the variance is low and the smoother is dominated by the incorrect observation. In the multi-view case, however, geometric consistency provides a powerful independent signal: if predictions across views cannot be explained by a single 3D point, at least one view must be wrong, regardless of how confident the ensemble is.

We exploit this signal through a procedure that inflates the observation noise for predictions that are geometrically inconsistent across views, down-weighting their influence on the posterior (Fig. 3d). When predictions are consistent and confident across views, the posterior recovers the true position without inflation (Fig. 3e). When predictions are inconsistent and the more confident view is correct, inflation downweights the incorrect view and the posterior recovers the true position (Fig. 3f). When the confident view is itself wrong, inflation cannot override it, but it correctly inflates the posterior uncertainty, flagging the frame as unreliable rather than propagating the error (Fig. 3g; Extended Data Fig. 4a,b; Supp Video 1).

This “variance-inflated” mvEKS outperforms non-inflated mvEKS across all datasets, with inflation concentrated predominantly on harder, more occluded keypoints, confirming that it detects genuine uncertainty rather than inflating indiscriminately (Extended Data Fig. 4c,d). The full progression LP3D →Ensemble Median → mvEKS shows consistent improvement at each stage across all datasets, with the nonlinear, variance-inflated version achieving the best performance on calibrated datasets (Fig. 3h). This pattern is illustrated clearly in the Fly dataset, where five keypoints per leg span from medial to distal joints, with distal keypoints being harder to track due to their smaller size and greater range of motion. LP3D improves over ResNet-50 across all joint positions, but the largest reductions in both 2D and 3D error occur at the most distal, challenging keypoints, and variance-inflated nonlinear mvEKS provides further improvements at each position (Extended Data Fig. 5).

We also compared mvEKS to Anipose (Karashchuk et al., 2021) on the Fly dataset. For a fair comparison, we applied Anipose to the ensemble median, the mvEKS input signal. Nonlinear mvEKS with variance inflation matches the performance of the full Anipose pipeline with 2D temporal filters and 3D spatiotemporal constraints (Extended Data Fig. 1d,e). However, mvEKS additionally provides posterior uncertainty estimates. Furthermore, mvEKS only requires a single automatically-selected hyperparameter per keypoint, whereas Anipose requires separate configuration of its 2D filtering and 3D constraint stages.

### 2.5 Uncertainty-guided pseudo-labeling closes the annotation gap for an end-to-end pipeline

Vision transformer performance scales with data size (Zhai et al., 2022), and LP3D inherits this property: accuracy continues to improve as more labeled frames are added (Extended Data Fig. 2). Yet manual annotation of additional frames remains a practical bottleneck for most experimental laboratories. A natural solution is to use an ensemble to generate pseudo-labels for unlabeled frames and distill its knowledge into a single model, eliminating the computational overhead of running multiple models at test time. However, this requires a principled way to identify which pseudo-labeled frames are trustworthy enough to use as supervision.

The uncertainty estimates produced by variance-inflated mvEKS resolve this tension. Because the posterior variance captures cross-view agreement rather than merely ensemble spread, frames where uncertainty is low across all keypoints and views can be trusted as high-quality pseudo-labels. We developed a simple method to retain a diverse set of these pseudo-labels to retrain a single LP3D model, transferring the knowledge of the full ensemble and mvEKS pipeline without requiring any additional manual annotation (Extended Data Fig. 6a; Methods). This process, known as distillation, trains a single model on the ensemble’s own predictions rather than on manual labels, producing a lightweight model that matches ensemble accuracy without the computational cost of running multiple models at inference time.

The distilled model outperforms individual LP3D ensemble members despite using identical architecture and training procedures, and when triangulation and reprojection are applied at inference, it matches the full LP3D + mvEKS pipeline at the cost of a single forward pass (Extended Data Fig. 6b). These gains are driven by the quality of frame selection, not the quantity of additional data: replacing uncertainty-guided selection with random selection leads to worse results (Extended Data Fig. 6c), confirming that the posterior variance is identifying genuinely informative frames rather than simply expanding the training set.

The result is a fully self-contained pipeline that requires no additional manual annotation and no dataset-specific tuning at any stage: train an LP3D ensemble on a small set of labels, apply mvEKS with variance inflation to generate high-quality pseudo-labels, and distill into a single efficient model. This directly addresses the annotation bottleneck that limits multi-view pose estimation in practice, while preserving the accuracy of the full ensemble pipeline at the cost of a single forward pass.

### 2.6 Lightning Pose 3D improves downstream neuro-behavioral analyses

Improved pose accuracy is only meaningful if it translates into better science. Noisy keypoint predictions introduce spurious kinematic features (e.g., erroneous inter-keypoint distances or speed transients) that contaminate downstream neuro-behavioral analyses. We used the IBL dataset (IBL et al., 2025) to evaluate whether LP3D’s improvements in pose accuracy produce measurable gains in downstream applications like unsupervised behavioral clustering and neural decoding.

To characterize the structure of paw movements derived from each model, we computed a range of kinematic features for each paw including 3D speed, velocity, acceleration and inter-paw distance (Fig. 4a). We clustered these features across *n* = 39 sessions and across ResNet-50 and LP3D + (linear) mvEKS predictions using *k*-means, and visualized the results with UMAP (McInnes et al., 2018); radar plots summarize the characteristic kinematic profile of each cluster (Fig. 4b,c). Hierarchical clustering on the shared feature space reveals a subset of clusters dominated by ResNet-50 predictions (clusters 2, 4, 5, 7; Fig. 4d). These clusters exhibit elevated per-paw reprojection errors, and coincide with occlusion events in which pose errors lead to spurious increases in paw speed, for example (Fig. 4d). In contrast, clusters containing LP3D + mvEKS predictions show lower reprojection errors and balanced kinematic feature profiles, reflecting genuine behavioral structure rather than tracking artifacts. Improved pose estimation therefore suppresses artifact-driven clusters and yields more stable, interpretable unsupervised behavioral representations.

**Figure 4:**
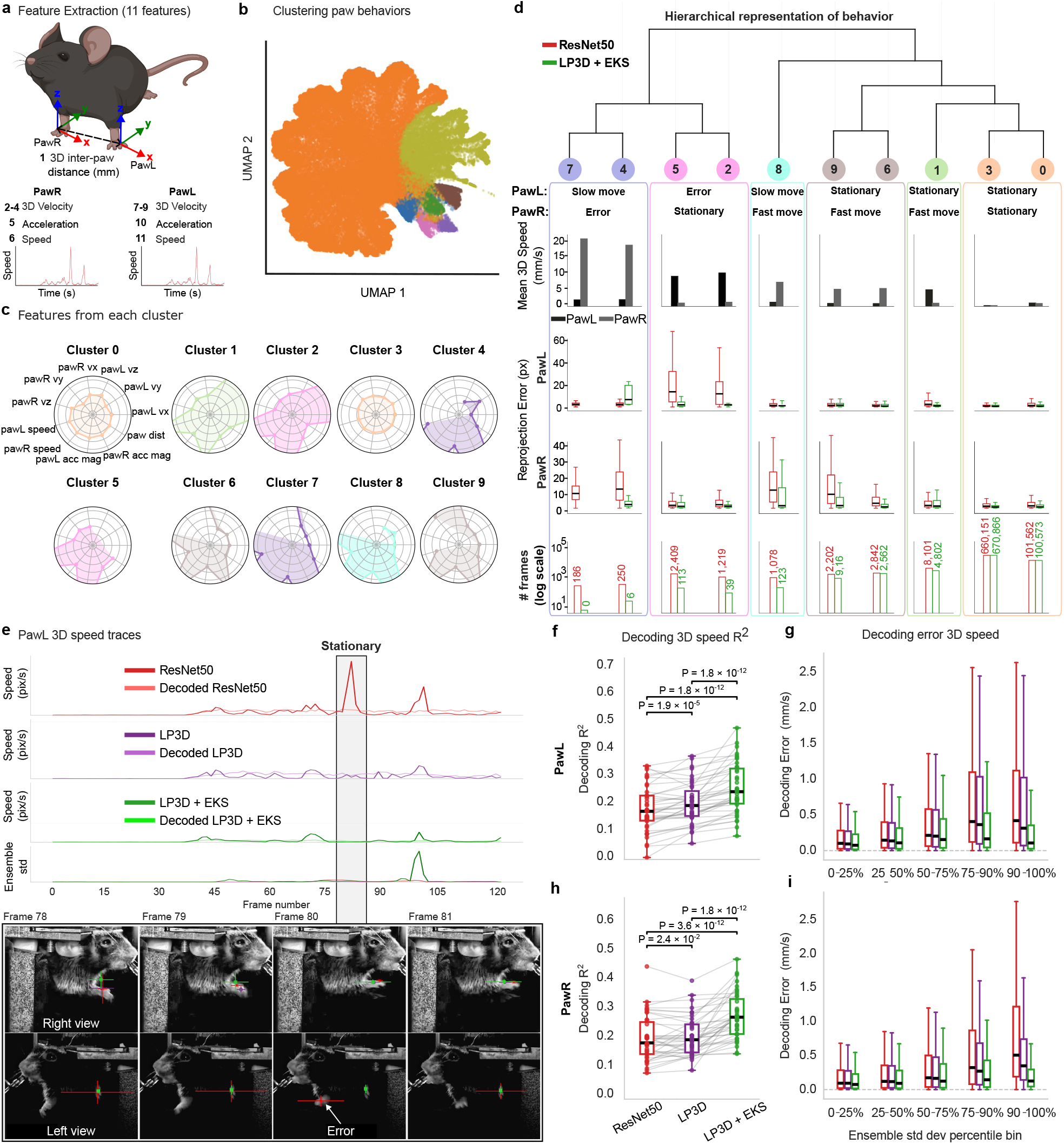
Accurate multi-view pose estimation improves downstream neuro-behavioral analyses in IBL dataset. **a**, Extracted behavioral features from paw: 3D speed, velocity and acceleration components for left and right paws, and inter-paw distance. **b**, UMAP embedding of kinematic features from a representative session, colored by *k*-means cluster. **c**, Radar plots of feature contributions per cluster. **d**, *Top*: dendrogram of cluster hierarchy. *Bottom*: mean 3D speed, reprojection error, and frame counts by model for each cluster. ResNet-50-dominated clusters (2, 4, 5, 7) show few LP3D+EKS frames and high reprojection error (occlusion artifacts). **e**, Left paw 3D speed—example trace (dark) and decoded (light)—for ResNet-50, LP3D, and LP3D+EKS; the gray shaded region marks a transient occlusion during which the paw is stationary. ResNet-50 exhibits a spurious speed spike within this window, while LP3D+EKS maintains a flat, accurate trace throughout. Image contrast enhanced to improve visibility. **f**, Decoding *R*^2^ for left paw 3D speed across *n* = 39 sessions; LP3D+EKS significantly outperforms ResNet-50 and LP3D (one-sided Wilcoxon signed-rank test). **g**, Left paw 3D speed decoding error (mm/s) as a function of ensemble standard deviation percentile; frames in larger percentile bins have larger uncertainty across models. Error increases with uncertainty for all models; LP3D+EKS shows lower error overall, with the gap widening in the 75–100% bins. **h**, Same as f, but for right paw 3D speed. i, Same as g, but for right paw 3D speed.

This result is illustrated concretely by a single occlusion event: ResNet-50 incorrectly localizes the paw during a transient occlusion, distorting the 3D speed trace, while LP3D + mvEKS maintains consistent tracking through the same event (Fig. 4e; Supp Video 2). This is precisely the failure mode that cross-view reasoning and variance inflation were designed to address.

These improvements translate directly into neural decoding accuracy (Fig. 4f-i; Table 2). We used cross-validated ridge regression to decode the 3D speed of both paws from multi-region Neuropixels recordings across *n* = 39 IBL sessions. LP3D + mvEKS achieves significantly higher decoding *R*^2^ for both paws compared to ResNet-50 and LP3D (one-sided Wilcoxon signed-rank test; Fig. 4f,h). To understand where these gains originate, we split frames into deciles based on pose prediction variance across all models, and computed decoding error within each bin. The performance gap between models widens substantially in the highest-variance deciles (Fig. 4g,i), where models disagree most and pose estimation is most challenging.

**Table 2:** Neural decoding performance in IBL data across pose estimation models. Values are mean ± SEM across *n* = 39 sessions. Decoding *R*^2^ is the coefficient of determination; decoding error is the mean absolute error (MAE) in mm/s.

| | | Decoding $R^2$ | Decoding error (mm/s) |
| --- | --- | --- | --- |
| IBL 3D paw speed (left) | ResNet-50 | 0.185±0.013 | 0.53±0.06 |
|  | LP3D | 0.208±0.012 | 0.51±0.06 |
|  | LP3D + mvEKS | 0.260±0.015 | 0.32±0.04 |
| IBL 3D paw speed (right) | ResNet-50 | 0.209±0.012 | 0.48±0.05 |
|  | LP3D | 0.216±0.011 | 0.41±0.04 |
|  | LP3D + mvEKS | 0.287±0.013 | 0.30±0.03 |

### 2.7 The Lightning Pose 3D software ecosystem

Methodological advances in pose estimation only have scientific impact if they are adopted by experimental laboratories. Yet most multi-view pipelines require users to assemble separate tools for calibration, labeling, training and post-processing, each with its own data formats and dependencies. This fragmentation prevents researchers from moving quickly from raw videos to initial pose estimates, and slows down the iterative cycle of inspecting predictions, labeling additional frames, and retraining that is essential for achieving high-quality results.

To address these issues, we designed the Lightning Pose 3D software ecosystem to be modular and accessible to users with a wide range of coding experience. The system is organized around a standardized project directory that contains labeled frames, videos and trained models, and can be accessed through three interfaces depending on the user’s needs: a command-line interface (CLI) that provides commands for training, inference, and other common operations; a Python API that exposes models directly (e.g., for use in Jupyter notebooks or custom scripts); and a browser-based user interface for interactive use (Fig. 5a). Because all three interfaces operate on the same project directory, users can freely combine them across deployment environments—local workstations, cloud machines, and HPC clusters—choosing the right interface and hardware for each task (Fig. 5b).

**Figure 5:**
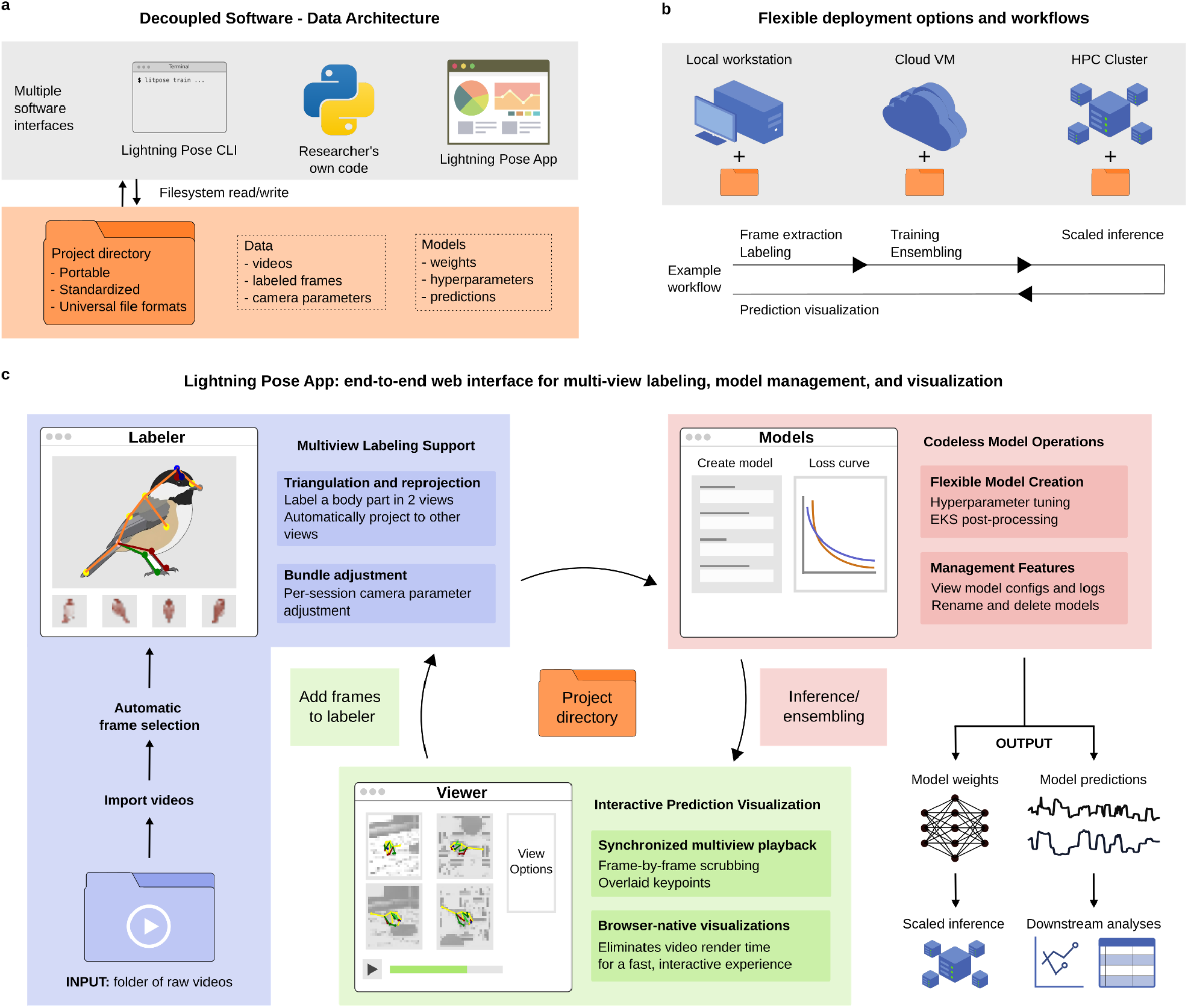
An easy-to-use interface for end-to-end multi-view pose estimation projects. **a**, The new app, existing CLI, and researcher’s own scripts all collaborate on a common “Project” via a uniform directory structure. **b**, The Project directory and software can be deployed on local workstations and headless servers, enabling various workflows. **c**, The modern web app is highly interactive and offers multi-view features: camera-calibration-assisted labeling and synchronized multi-view video playback, overlaid prediction markers and real-time scrubbing. The modules are tightly integrated allowing a closed loop from labeling to model fine-tuning to prediction viewing and back to labeling.

To make these tools widely accessible, we developed a browser-based application that supports both single- and multi-view pose estimation projects (Fig. 5c). The labeling module supports camera-parameter-assisted annotation, inspired by the Anivia labeler (Karaschuk, 2026): a user labels a body part in as few as two views, and the app triangulates and reprojects the label to all remaining views, significantly reducing annotation time for setups with many cameras (Maree et al., 2024). In the model training module, users can select from previous single-view Lightning Pose models (Biderman et al., 2024) as well as the SVT and MVT architectures presented here, and have access to a set of hyperparameters with sensible defaults directly in the app. After training, inference and EKS post-processing can be launched with a single click. The prediction viewer module allows users to scrub through videos with model predictions overlaid; frames can be selected and sent directly to the labeling module, where model predictions serve as initial labels that the user can correct. This closed feedback loop, from prediction inspection to targeted labeling to retraining, allows users to efficiently identify and correct failure cases without leaving the interface.

## 3 Discussion

We introduced Lightning Pose 3D (LP3D), a flexible end-to-end framework for multi-view animal pose estimation that combines a data-efficient multi-view transformer architecture, novel training augmentations and losses, an updated Ensemble Kalman Smoother (EKS) post-processor with better-calibrated uncertainty estimates, and an uncertainty-guided pseudo-labeling pipeline. An accompanying browser-based user interface supports all stages of a multi-view pose estimation project.

When designing LP3D, we prioritized flexibility, generalizability, and usability. The framework flexibly operates across a range of experimental conditions: the multi-view transformer (MVT) and linear EKS function without camera calibration, while 3D augmentations, reprojection loss, and nonlinear EKS leverage calibration information when available. The MVT architecture is agnostic to the choice of vision transformer backbone; we use a DINO-pretrained ViT-S in this work (Caron et al., 2021), but the same framework accommodates other pretrained vision transformers (He et al., 2022; Assran et al., 2023; Wang et al., 2025; Wang et al., 2026), allowing LP3D to incorporate stronger backbones as they become available. To demonstrate generalizability, we validated LP3D on five datasets spanning three species, including setups ranging from two to six cameras, and a multi-animal dataset with two visually distinct individuals. We use a single set of training hyperparameters across all datasets (Table 1) and show robustness to cross-session camera drift (Supp Fig. 10). To support end-to-end usability, we developed a browser-based application for frame extraction, multi-view labeling, model training and inference, post-processing, and diagnostic visualization (Fig. 5). A consistent user interface is a crucial counterpart to the methodological flexibility highlighted above: as underlying models and best practices evolve, users can adopt improvements without needing to engage with low-level technical details or rebuild their workflows.

The most common approach to multi-view pose estimation processes each camera view independently with a 2D pose network and fuses predictions post hoc via triangulation (Nath et al., 2019; Karashchuk et al., 2021). We have shown that this view-independent design is consistently outperformed by methods that incorporate multi-view information during feature extraction, particularly under occlusion and with limited labeled data (Fig. 2). Among methods that do process views jointly, DANNCE (Dunn et al., 2021) is perhaps the most widely known in the behavioral neuroscience community. DANNCE constructs a learned volumetric representation from all views, and while effective, this approach requires precise camera calibration, limiting its applicability. LP3D, by contrast, operates with or without camera parameters. In addition to improved accuracy (Extended Data Fig. 1), LP3D’s non-volumetric representations lead to ∼15-20% faster inference times (Supp Fig. 2). Additionally, LP3D’s backbone-agnostic design allows it to leverage pretrained vision transformers, which contributes to its strong performance with limited labeled data.

The transition from single-to multi-view pose estimation introduces a wealth of geometric structure that is absent in the single-view setting. In this work we focused on fully exploiting this structure on labeled data through the multi-view transformer, patch masking and the 3D reprojection loss, which together already provide large gains over independent-view baselines (Fig. 2). The original Lightning Pose framework (Biderman et al., 2024) took a complementary approach, introducing unsupervised losses (including multi-view and pose consistency losses) that enabled training on unlabeled frames. In the current LP3D framework, the 3D reprojection loss applies the multi-view consistency principle to labeled data; extending both multi-view and 3D pose consistency losses (Extended Data Fig. 7) to both labeled and unlabeled data is a concrete next step that would allow the model to train on substantially larger datasets without additional annotation cost.

The pipeline presented here handles single animals and multiple visually distinct animals. Extending it to the most general case of multiple visually indistinct animals represents an important direction for future work. We have already demonstrated that LP3D achieves high-quality pose estimation within a cropped bounding box on the chickadee dataset, following a top-down approach (Pereira et al., 2022). To extend this to multiple animals, we would first need to estimate bounding boxes and assign identities in each view. Computer vision foundation models are becoming increasingly capable at these tasks (Yang et al., 2026; Carion et al., 2025), and we expect rapid continued progress. Once identities and bounding boxes have been matched across views a single-animal pose estimator, like those presented here, can be deployed within each cropped region (Klibaite et al., 2025). We believe this modular approach, which offloads detection and identity tracking to rapidly improving foundation models and reserves LP3D for high-precision pose estimation, is preferable to combining these tasks into a single model.

As vision-language models (VLMs) become more capable (Zhang et al., 2024), they offer a path toward further reducing annotation cost (Ye et al., 2023; Ke et al., 2026). A user could provide a list of keypoint names and optional context, and a VLM could propose initial keypoint locations across a large number of frames. The multi-view and pose consistency metrics we have introduced provide natural quality checks: frames with high inconsistency can be flagged for human review and correction. Such a semi-automated annotation process would complement the advances in network training and post-processing presented here, further lowering the barrier to high-quality multi-view pose estimation. More broadly, this modular approach to incorporating new capabilities will allow LP3D to continue delivering state-of-the-art performance as the field of computer vision advances, helping researchers address pressing scientific questions about animal behavior.

## Acknowledgments

We thank Peter Dayan and Anne Churchland for serving on our IBL paper review board; Farzad Ziaie for help with early versions of the multi-view data loading code; Lili Karaschuk for discussions about multi-view labeling; Cole Hurwitz for discussions of transformers and multi-view EKS; Yiting Chang, Sherry Li, Jennifer Mejaes, Catalin Mitelut, Simon Nilsson, the Zirong Lab, and Jessica Zung for testing out early versions of the pipeline and app; and Lior Aharon for help with writing and figures. This work was supported by the following grants: Gatsby Charitable Foundation GAT3708 (LA, MRW, KS, KL, YW, LP), NIH DP1-MH136573 (IW), NIH F32MH123015 (SC), NIH K99NS136846 (SC), NIH 1R50NS145433 (MRW), NIH U19NS123716 (MRW, KS, IBL, LP), NSF 1707398 (LA, MRW, KS, KL, YW, LP), the National Science Foundation and DoD OUSD (R&E) under Cooperative Agreement PHY-2229929 (The NSF AI Institute for Artificial and Natural Intelligence) (LP), Simons Foundation 543023 (MRW, IBL, LP), Wellcome Trust 216324 (IBL), and funds from Zuckerman Institute Team Science (MRW). The funders had no role in study design, data collection and analysis, decision to publish or preparation of the manuscript.

## Author Contributions Statement

Conceptualization: LA, MRW, LP; Software package development: LA, MRW, KS, KL; User interface development: KS; Data collection: BM, SC; Data analysis: LA, YW; First draft (writing): LA, MRW, LP; First draft (editing): LA, MRW, LP; Funding: IW, DA, LP.

## 4 Methods

### 4.1 Datasets

All datasets used for the paper were collected in compliance with the relevant ethical regulations. See the following published papers for each dataset: Treadmill Mouse (Warren et al., 2021), Fly (Karashchuk et al., 2021), Chickadee (Chettih et al., 2024), IBL (IBL et al., 2025a), and Rat7M (Marshall et al., 2021). See below for more information on the Resident-Intruder dataset.

#### Treadmill mouse

Head-fixed mice run on a circular treadmill while avoiding a moving obstacle (Warren et al., 2021). The treadmill has a transparent floor and a mirror mounted inside at 45°, allowing a single camera to capture two roughly orthogonal views (side and bottom views via the mirror) at 250 Hz. Frame sizes are 406 × 396 pixels. We split each frame vertically into its respective views in order to make a “multicamera” dataset. Each view is reshaped during training to 128 × 256 pixels. Seven keypoints on the mouse’s body are labeled in each view. The training/test sets consist of 789/253 instances, respectively. We use a 3D latent space for linear mvEKS (Extended Data Fig. 3c). We accessed the labeled pose estimation dataset from https://doi.org/10.6084/m9.figshare.24993315.v1 under the CC-BY 4.0 license.

#### Fly

Head-fixed flies behave spontaneously on an air-supported ball, captured by six cameras at 300 Hz (Karashchuk et al., 2021). Frame sizes vary by view, but are ∼450×525, and frames are reshaped during training to 256×256 pixels. Thirty keypoints are labeled in each view: five joints on each of six legs.

Our pose estimation models require labels for all views at a given instant in time, and although some of this data is available in the Anipose repository (https://datadryad.org/dataset/doi:10.5061/dryad.nzs7h44s4), we took a different approach to ensure a large quantity of high-quality, simultaneously labeled frames. For a subset of sessions in the data repository that contain Anipose predictions, we treat a subset of these predictions as labels for training our own models. We first remove any instance where average 3D reprojection error is *>*10 pixels. We then run *k*-means clustering on the remaining 3D poses (using 25 clusters per session) and select one example per cluster. The training/test sets consist of 377/300 instances, respectively. We use a 3D latent space for linear mvEKS (Extended Data Fig. 3c).

#### Chickadee

Freely moving chickadees engage in seed caching behavior in a large arena, captured by six cameras at 60 Hz (Chettih et al., 2024). Frame sizes vary by view but are approximately 3000 × 1500 pixels. We created a cropped dataset using the ground truth labels to define a bounding box around the bird, and reshaped the cropped frames to 320 × 320 pixels. Each view is reshaped during training to 256 × 256 pixels. Eighteen keypoints on the chickadee’s body are labeled in each view. The training/test set consists of 433/143 instances, respectively. We use a 6D latent space for linear mvEKS (Extended Data Fig. 3c).

To produce the cropped unlabeled videos for distillation, we implemented a two-stage top-down pose estimation pipeline (Pereira et al., 2022). First, we trained a detector network on full resolution frames down-sampled to 256 × 256 pixels to localize the bird within each frame. We then computed a bounding box around the bird in each view, ran inference using a pose estimation model trained specifically on cropped frames, and transformed the resulting predictions back to full-resolution coordinates before applying mvEKS.

#### IBL

Head-fixed mice perform a visually-guided decision-making task (IBL, 2023; IBL et al., 2025b; IBL et al., 2025a; Findling et al., 2025). Two cameras—’left’ (60 Hz, 1280 × 1024 pixels) and ‘right’ (150 Hz, 640 × 512 pixels)—capture roughly orthogonal side views of the mouse’s face and upper trunk during each session (IBL et al., 2022). We created a temporally downsampled version of the right video using nearest neighbor interpolation between the left and right side video timestamps. We further spatially downsampled each view to 320 × 256 pixels. We used the Lightning Pose 3D app to extract and label frames. We tracked three keypoints per view, one for each paw and one for the nose. The training/test sets consist of 640/280 instances, respectively. We use a 3D latent space for linear mvEKS (Extended Data Fig. 3c).

#### Resident-Intruder

Two male mice (a C57/Bl6 resident and a Swiss-Webster intruder; Jackson Laboratory and Taconic Biosciences, respectively) interact freely for five minutes per day as part of a chronic social stress experiment. Five cameras (Flir Blackfly S USB, 6mm Fujinon lenses) are positioned at various angles above and beside a transparent plexiglass cage to maximize coverage of the cage interior. Videos are acquired at 120 Hz at 1280 × 1024 pixels under IR illumination, with acquisition synchronized across cameras via TTL pulses from an Arduino Nano. All experimental procedures were approved by the Princeton University Institutional Animal Care and Use Committee (Protocol #1876). Twenty-two keypoints are labeled per frame: 11 per animal. The training/test sets consist of 207/38 instances, respectively. We use a 3D latent space for linear mvEKS (Extended Data Fig. 3c).

#### Rat7M

A single freely moving rat behaves spontaneously in an open field, captured by six cameras at 120 Hz (Marshall et al., 2021). Frame sizes are 1328 × 1048 pixels. We create a cropped dataset using the ground truth labels to define a bounding box around the rat, and reshaped the cropped frames to 320 × 320 pixels. Each view is reshaped during training to 256 × 256 pixels. We retain 15 of the original keypoints in each view, omitting those tracking the head stage of the rat.

Rather than retain the 30,000+ labeled instances in the original dataset, we chose to curate a smaller subset of instances matched in size with our other datasets. Similar to the Fly dataset, for each session we first filter out any instances with missing data. We then filter out potentially problematic points using skeleton distances (`ElbowR`-`ElbowL`distance in [40, 60]; `KneeR`-`ShinR`and `KneeL`-`ShinL`distances in [10, 1000]). We then run *k*-means clustering on the remaining 3D poses (using 100 clusters per session) and select one example per cluster. Finally, we performed manual inspection of the resulting labels and excluded any instances where ground truth keypoints were incorrect due to camera syncing issues. The training/test sets consist of 455/177 instances, respectively. For the results in Extended Data Fig. 1 we only compute error on 96 test frames that the original DANNCE model was not trained on. We accessed the videos and motion capture labels from https://doi.org/10.6084/m9.figshare.c.5295370 under the CC-BY 4.0 license.

### 4.2 Model architectures

#### Single-view baselines

Our single-view models perform heatmap regression on a frame-by-frame basis, akin to DeepLabCut (Mathis et al., 2018), SLEAP (Pereira et al., 2022), DeepPoseKit (Graving et al., 2019), and others. Each has roughly the same architecture: a “backbone” network that extracts a set of 2D feature maps per frame, and a “head” that transforms these into a predicted heatmap for each labeled keypoint. The head includes a series of identical ConvTranspose2D layers that each double the heatmap in size; see Biderman et al., 2024 for more details. Each heatmap is normalized with a 2D spatial softmax with a temperature parameter *τ* = 1. The supervised loss is the mean square error between predicted heatmaps and labeled heatmaps, averaged over all keypoints and batch elements.

Once heatmaps have been predicted for each keypoint, we transform these 2D arrays into estimates of the width-height coordinates in the original image space. We first upsample each heatmap using bicubic interpolation. A 2D spatial softmax renormalizes the heatmap to sum to 1, and we apply a high temperature parameter (*τ* = 1000) to suppress non-global maxima. A 2D spatial expectation then produces a subpixel estimate of the location of the heatmap’s maximum value. These two operations – spatial softmax followed by spatial expectation – are together known as a soft argmax (Wu et al., 2020). To compute the confidence value associated with the pixel coordinates, we sum the values of the normalized heatmap within a configurable radius of the soft argmax.

We compared a variety of vision transformer backbones easily accessible through Hugging Face:

- ViT/B-16 pretrained on ImageNet with masked autoencoding (He et al., 2022), available at https://huggingface.co/facebook/vit-mae-base. The “base” ViT contains ∼80M parameters, which is 4x larger than the ResNet-50 (∼20M parameters). The “16” indicates the model utilizes a patch size of 16×16 pixels.
- ViT/B-16 pretrained on ImageNet with DINO (Caron et al., 2021), available at https://huggingface.co/facebook/dino-vitb16.
- ViT/S-16 pretrained on ImageNet with DINO, available at https://huggingface.co/facebook/dino-vits16. The small ViT contains ∼20M parameters, on par with ResNet-50.
- ViT/B-16 pretrained on multiple datasets with DINOv2 (Oquab et al., 2023), available at https://huggingface.co/facebook/dinov2-base.
- ViT/S-16 pretrained on multiple datasets with DINOv2, available at https://huggingface.co/facebook/dinov2-small.

We train models for each backbone with three random seeds (see below) and compare to our ResNet-50 baseline pretrained with the AnimalPose10K dataset (Yu et al., 2021). The transformers outperform the ResNet across all datasets except for Fly, though no single backbone consistently performs best across datasets (Supp Fig. 1a). Given these results, we chose V_I_T/S-DINO for our subsequent experiments due to considerably faster training time than the “base” models (2-3× faster; Supp Fig. 2) and lower memory requirements, an important constraint for our domain application where we expect individual labs to be running these models on single consumer-grade GPUs.

#### Multi-view transformer (MVT)

Many multi-view pose estimation techniques employ bespoke architectural elements. While these architectures may provide good performance with enough training data, they do not allow us to easily exploit general pretrained backbones that are useful when training models with a small number of labels. Furthermore, algorithmic simplicity is desirable for our application domain, where users are often experimental labs with little experience maintaining and debugging exotic architectures. Here we propose a simple strategy that allows the model to take advantage of multiple views and is also compatible with generic vision transformer (V_I_T) backbones: rather than process pixel patches from each view independently, we process all patches simultaneously, allowing the standard self-attention mechanism to pool information within and across views.

The standard image V_I_T (Dosovitskiy et al., 2020) data pipeline starts with a 2D image **x** ∈ ℝ^*H*×*W* ×*C*^ (where *H, W, C* are height, width, channels) and splits it into 2D patches, each with shape (*P* × *P* × *C*), where the patch size *P* is typically 16. Each patch is reshaped to a vector of length *P* ^2^*C*, and all patches are concatenated into a sequence of the *N* flattened 2D patches 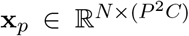, where *N* is the total number of patches. Each flattened patch **x**_*p,i*_ is mapped with a trainable linear projection to a patch token 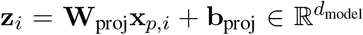, where *d*_model_ is the embedding dimension. A standard fixed 1D position encoding 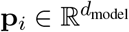 is added to the patch tokens to retain information about the patch location.

To extend this framework to *V* camera views (Fig. 1f), we apply the same patch embedding pipeline independently to each view **x**^(*v*)^, *v* = 1, …, *V*. Each patch is projected and enriched with a fixed positional encoding **p**_*i*_ as before, and an additional learnable view encoding **v**_*v*_ (inspired by (Ma et al., 2021; Zhou et al., 2023)), resulting in 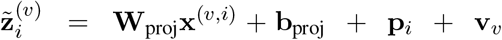. Concatenating all patches from all views forms the joint input sequence 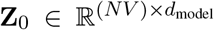 for the V_I_T encoder, which produces **Z**_Enc_ = ViTEncoder(**Z**_0_). We regroup **Z**_Enc_ by view and reshape to the original 2D patch grids. Following ViTPose (Xu et al., 2022), which showed that V_I_T backbones retain accuracy with minimal decoders, we employ the lightweight shared upsampling head described above to map each per-view grid into keypoint heatmaps.

#### MVT backbones

The power of our multi-view transformer approach is that it does not require any bespoke or complex architectures, which can require large amounts of data to properly train (Nogueira et al., 2025). Instead, we use encoders from off-the-shelf pretrained transformers combined with simple heatmap heads, which (1) reduces the number of parameters we need to train from scratch; and (2) forces all of the complex cross-view information propagation into the backbone. We tested the DINO-based vision transformer backbones introduced above, and find that performance is dataset-dependent (Supp Fig. 1b); therefore we again choose V_I_T-S/DINO as our default MVT backbone due to its good speed-accuracy tradeoffs.

### 4.3 Pose estimation training

#### Patch masking

The success of masked autoencoding in self-supervised vision transformers (He et al., 2022) inspired us to take a similar approach in the supervised domain of pose estimation. For each labeled instance, we randomly select patches and zero their pixel values before adding position and view encodings. This data augmentation mimics occlusions and forces the transformer to fully exploit cross-view self-attention. We use curriculum learning starting after 700 iterations with 10% patch masking per view, linearly increasing to 50% by iteration 5000.

#### 2D augmentations

We closely follow the 2D augmentation pipeline of DeepLabCut (Mathis et al., 2018). We apply the same non-geometric augmentations to all datasets and models using the `imaug`package with the indicated parameters and probabilities *p*:

- `MotionBlur(k=5, angle=90)`, *p* = 0.5
- `CoarseDropout(p=0.02, size_percent=0.3, per_channel=0.5)`, *p* = 0.5
- `CoarseSalt(p=0.01, size_percent=(0.05, 0.1))`, *p* = 0.5
- `CoarsePepper(p=0.01, size_percent=(0.05, 0.1))`, *p* = 0.5
- `AllChannelsHistogramEqualization()`, *p* = 0.1
- `Emboss(alpha=(0, 0.5), strength=(0.5, 1.5))`, *p* = 0.1

For the single-view models (both ResNets and transformers), we apply additional geometric augmentations, i.e. those which affect the locations of the corresponding keypoints:

- `Affine(rotation=(-25, 25))`, *p* = 0.4
- `ElasticTransformation(alpha=(0, 10), sigma=5)`, *p* = 0.5
- `CropAndPad(percent=(-0.15, 0.15), keep_size=False)`, *p* = 0.4

#### 3D augmentations

Standard data augmentation pipelines apply transformations independently to each view, creating problems for multi-view models–particularly those using a 3D consistency loss–that require geometrically consistent augmentations across views for each labeled instance. We therefore implement a 3D data augmentation scheme that maintains geometric consistency, while still applying 2D non-geometric augmentations independently to each view.

First, we triangulate 2D ground truth labels using camera parameters to obtain 3D keypoint positions. We randomly scale keypoints by median-centering, multiplying by a random factor drawn from 𝒰 (0.8, 1.2), then reapplying the median. Next, we randomly translate keypoints by computing a bounding box in each dimension using the minimum and maximum keypoint coordinates, multiplying its width by a random factor from 𝒰 (− 0.25, 0.25), and shifting keypoints by the result (such that the shift will be a maximum of 25% of the width of the animal in any direction). Since camera parameters remain fixed, this is equivalent to scaling and translating the subject within the recorded area. To augment images, we reproject the transformed 3D keypoints back to each camera view, estimate view-specific affine transformations between original and augmented labels using OpenCV’s `estimateAffinePartial2D`, then apply these transformations to the images (Supp Fig. 4).

#### 3D reprojection loss

We use predicted heatmaps and camera calibration parameters to enforce multi-view geometric consistency independently across keypoints. We compute the soft argmax (2D spatial expectation) of predicted heatmaps for each view to extract 2D coordinates. This differentiable operation enables downstream losses to use these coordinate estimates. Using the ground truth camera parameters, we triangulate these predicted 2D coordinates across all camera pairs via OpenCV’s `undistortPoints`and `triangulatePoints`, following Karashchuk et al., 2021, yielding a set of per-view-pair 3D estimates. We take the mean across all camera pairs to obtain a single 3D position per keypoint.

We then reproject these mean 3D positions back into 2D for each camera view using the same camera parameters. This forces every view to incorporate information from all other views. From the reprojected 2D coordinates, we generate Gaussian heatmaps and compute mean squared error against the ground truth heatmaps, scaling by the number of heatmap pixels (*h* × *w*) to standardize the loss range. Missing keypoints (e.g., those that moved outside the frame boundary during augmentation) are excluded from the computation. The final loss is the mean square error across all valid keypoints and views in the batch, weighted by a hyperparameter that balances this term with the 2D heatmap loss. Importantly, this loss is on the same scale as the original 2D heatmap loss, simplifying hyperparameter tuning. We find the same weight works well across all datasets (`log_weight=3`in our configuration file, corresponding to a weight of 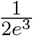) and therefore use this value for all experiments (Supp Fig. 6). The weight on the 3D projection loss is initially set to 0 for the first 60 epochs (so the iteration number is dataset-dependent), and linearly anneal the weight from 0 to its full value over the remaining iterations.

#### Training details

We use Lightning Pose (Biderman et al., 2024) to train supervised single-view pose estimation models on each dataset, and the extensions detailed above for multi-view pose estimation. For training and inference in both settings, we load all camera views simultaneously for each time point, such that each batch element comprises one image per camera view (e.g., with six views, a batch size of four contains 24 total images). We use a batch size of eight instances per network. During training of single-view models, we apply standard image augmentations to labeled frames, including geometric transformations (rotations, crops), color space manipulations (histogram equalization), and kernel filters (motion blur). For data augmentation in multi-view models with camera calibration, see above.

We split the non-test data into 95% for training and 5% for validation. To simulate a limited-data scenario, we randomly select only 100, 200, or `min(400, total_train_frames)`instances from the training set. All evaluations use the model iteration with the lowest validation loss. Different ensemble members use different random seeds for the train/validation split.

We train models for 5000 iterations using the Adam optimizer (Kingma et al., 2014) with an initial learning rate of 1*e*-3 for the ResNet backbones and 5*e*-5 for the vision transformer backbones, which is halved at iterations 2000, 3000 and 4000. The pretrained backbone remains frozen for the first 400 iterations.

### 4.4 MVT data scaling

All analyses use models trained with 200 labeled frames unless otherwise noted, as this represents a reasonable annotation effort for a given experimental setup. To understand how each contribution — multi-view transformer, patch masking, and 3D loss — scales with data, we also train models with 100 and 400 labeled frames (Extended Data Fig. 2). For each combination of backbone and label count, we train three models with different random seeds. Labeled frame subsets are nested across counts within each seed, so that the 100 frames are a subset of the 200, which are a subset of the 400.

### 4.5 DeepLabCut baseline

For DeepLabCut (Mathis et al., 2018) experiments (Extended Data Fig. 1a), we use the default configuration: an ImageNet-pretrained ResNet-50 backbone trained for 300 epochs with the Adam optimizer (Kingma et al., 2014) and a learning rate of 1*e*-3, which matches the learning rate used for the Lightning Pose ResNet-50 models. The batch size was set to 8× the number of views to match the effective batch size of the Lightning Pose models. For each dataset, we trained three models on 200 labeled frames with different training/validation splits, using the same splits as the corresponding Lightning Pose models in all analyses.

### 4.6 DANNCE baseline

For DANNCE (Dunn et al., 2021) experiments, we used the Social DANNCE (SDANNCE) codebase^1^.

#### Accuracy

To evaluate pretrained performance, we applied the `DANNCE_comp_pretrained_rat7m.pth`backbone, which was trained on 30,076 Rat7M frames^2^, to run inference on 96 test frames from Rat7M, and compared against a single LP3D network trained on 455 labeled frames (Extended Data Fig. 1b).

For the Fly and Chickadee datasets, we trained DANNCE models both from scratch and by fine-tuning from the `DANNCE_comp_pretrained_single+r7m.pth`backbone, following the recommended fine-tuning procedure in the SDANNCE documentation^3^ and demo training script^4^. We report the better-performing variant for each dataset: fine-tuning for Fly and training from scratch for Chickadee. All models used a compressed DANNCE architecture (`net_type=compressed_dannce`) in mono mode with L1 loss, a learning rate of 1 × 10^−4^, batch size of 4, and 2,000 epochs. The 3D volume was defined over a 64^3^ voxel grid with nearest-neighbor interpolation and expected-value decoding (`expval=true`); volume bounds were set to ±42 mm for Chickadee (following Dunn et al., 2021 guidelines) and ±1.5 mm for Fly, chosen so that the voxel grid fully enclosed the animal, as verified by visual inspection. Random view replacement was enabled with all six views sampled per training example (`n_rand_views=6`); hue and mirror augmentations were disabled. Centers of mass were computed from labels (`com_fromlabels=true`). For each dataset, we trained three models on 200 labeled frames using the same training/validation splits as the corresponding LP3D models (Extended Data Fig. 1c).

#### Timing

We evaluated the computational performance of LP3D and DANNCE on the chickadee dataset using a batch size of 16 on an NVIDIA L4 GPU, reporting the mean ± s.e.m. over 100 batches. LP3D achieved faster training throughput than DANNCE (2.22 ± 0.02 vs. 2.81 ± 0.01 s/batch), and similarly outperformed DANNCE at inference (23.44 ± 0.39 vs. 20.68 ± 0.24 FPS), demonstrating that the improved accuracy of LP3D does not come at a computational cost relative to the baseline.

### 4.7 Anipose baseline

To produce 3D pose estimates from multi-view 2D predictions, we ran the Anipose triangulation pipeline (Karashchuk et al., 2021) (`anipose`v1.1.24, `aniposelib`v0.7.2) with its published default parameters. For the Fly dataset, 2D predictions from each of the six cameras were first processed through two sequential 2D filters: (1) a Viterbi-based temporal filter (`n_back=5`, `offset_threshold=25`, `score_threshold=0.05`; spline interpolation enabled), which resolves identity swaps and removes low-confidence outliers per key-point track, and (2) a points-based autoencoder filter, which learns the expected spatial layout of all 30 leg keypoints from labeled data via an MLP regressor trained with noise augmentation, and replaces 2D coordinates whose adjusted score falls below 0.3 with the autoencoder’s prediction. This two-stage 2D filtering is identical to the default Anipose processing pipeline.

Following 2D filtering, Anipose’s triangulation routine discards 2D detections whose likelihood score falls below a default threshold of 0.8 (setting the corresponding rays to NaN before triangulation). We retained this default behavior, evaluating the threshold against post-Viterbi scores rather than post-autoencoder scores, since the latter are on a different scale than DeepLabCut/Lightning Pose likelihoods. For 3D reconstruction, an initial estimate was obtained via direct linear transformation (DLT) triangulation, after which Anipose’s constrained optimization (`optim_points`) was applied. We evaluated two method variants:

- **2D-filters only**: Viterbi + autoencoder filtering without triangulation, providing a 2D baseline.
- **Spatiotemporal**: DLT triangulation followed by constrained optimization combining temporal smoothness (`scale_smooth=2`, `n_deriv_smooth=3`) and bone-length constraints (`scale_length=2`, `scale_length_weak=1`, with the skeleton defined by 24 segment pairs across six legs).

The triangulation used a reprojection error threshold of 5 pixels and a soft-ℓ_1_ reprojection loss, matching Anipose’s defaults.

### 4.8 Linear Ensemble Kalman Smoother

We first discuss the PCA-based linear version of mvEKS, describing both parameter initialization and our new automatic smoothing procedure. This version of mvEKS relies on the low intrinsic dimensionality of the multi-view data, which we find holds across all datasets (Extended Data Fig. 3c), and does not require camera calibration. The next section describes the nonlinear mvEKS, which can provide improved performance, especially for setups with larger lens distortion.

The linear EKS model for a single body part described in the main text is:

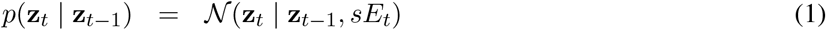

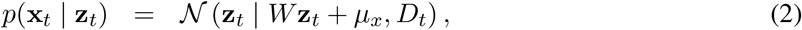

where **x**_*t*_ represents the (*x, y*) coordinates of a given body part across all views at time *t*, and **z**_*t*_ represents the underlying 3D (*x, y, z*) coordinates at *t*. We initialize model parameters by restricting to frames with low ensemble variance and use Principal Component Analysis (PCA) to estimate *W* and *µ*_*x*_. We then take temporal differences of the resulting PCA projections and compute their covariance to initialize *E*_*t*_. Finally, we set *D*_*t*_ as a diagonal matrix defined by the ensemble variance at time *t*.

Selecting the optimal smoothing parameter *s* in Eq. 1 is crucial: too large leads to undersmoothing, while too small causes oversmoothing (Fig. 3c). The optimal parameter occupies a well-defined minimum in the log-likelihood loss landscape (Extended Data Fig. 3a) and must be learned from data, as it varies substantially across keypoints and videos (Extended Data Fig. 3b). To perform automatic tuning of this parameter, we use the Adam optimizer implemented in the `optax`package with learning rate set to 0.25.

We initialize *s* using the standard deviation of the temporal differences of the initial PCA projections, which we have found to often lie near the minimum of the log-likelihood loss function. To improve the computational efficiency, we implemented a version of EKS that parallelizes over keypoints using the `JAX`library (Bradbury et al., 2018).

### 4.9 Nonlinear Ensemble Kalman Smoother

The nonlinear mvEKS model, again applied one body part at a time, uses camera calibration parameters to map the latent state **z**_*t*_ in Eq. 1 to the observations 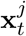 in view *j*:

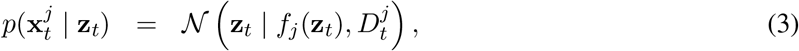

where *f*_*j*_ represents a standard camera model with radial and tangential distortion (Hartley et al., 2003).

For Fly, Chickadee, IBL, and Resident-Intruder datasets these parameters are obtained using standard camera calibration techniques, as described in Karashchuk et al., 2021.

We implement the nonlinear Kalman filter-smoother recursions using the `Dynamax`library (Linderman et al., 2025).

### 4.10 Variance inflation for the Ensemble Kalman Smoother

The EKS models above assume that ensemble variance *D*_*t*_ reliably captures prediction uncertainty. When all ensemble members agree on an incorrect prediction, however, this assumption breaks down, and the low variance causes the smoother to trust the erroneous observation (Fig. 3g). Multi-view geometry offers a remedy: predictions that cannot be reconciled into a single 3D point reveal that at least one view is incorrect, regardless of ensemble confidence (Fig. 3d). Our variance inflation procedure uses this signal to identify geometrically inconsistent views and inflate their observation noise *D*_*t*_, reducing their influence on the posterior. Here we derive the inconsistency metric underlying this procedure, based on a “Mahalanobis distance” in a linear latent variable model (Bishop, 2006), that quantifies whether each view’s prediction for a given body part is consistent with the remaining views.

We start with the general case of computing the Mahalanobis distance with an uninformative prior, which will be used for each body part at each point in time across all views; the next section details the application of this metric to multi-view pose estimation for outlier detection.

Let **x** ∈ ℝ^*n*^ be a vector of observations. We describe these observations with a standard linear latent variable model:

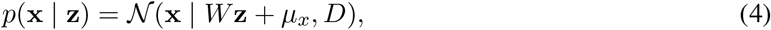

where **z** ∈ ℝ^*d*^ is a set of unobserved latent variables. For simplicity we will assume *D* is a diagonal matrix (note that with this assumption Eq. 4 becomes the generative model of Factor Analysis (Bishop, 2006)).

The posterior distribution of the latents given the observations is

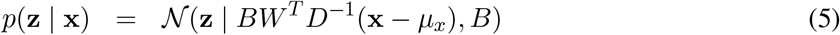

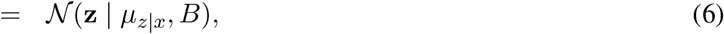

where *B* = (*I* + *W*^*T*^ *D*^−1^*W*)^−1^ if the prior on **z** is 𝒩 (0, *I*). However, this is a strong assumption, and instead we can use an uninformative prior where **z** ∼ lim_*σ*→∞_ 𝒩 (0, *σI*); this results in

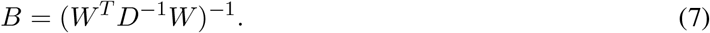

Next we will consider the posterior predictive distribution *p*(**x**^′^ | **x**), which describes the distribution of a new observation **x**^′^ given the observed data **x**:

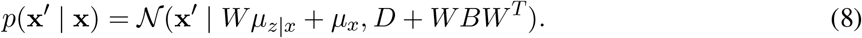

We can now use this information to compute the distance between the original observation **x** and the posterior predictive distribution, which is essentially measuring the reprojection error of the observation scaled by a covariance matrix. If we define *Q* = *D* +*WBW*^*T*^ (the covariance of the posterior predictive distribution), then the Mahalanobis distance is computed as

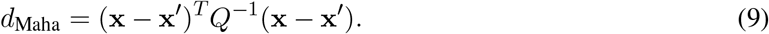

For more information on these derivations see Bishop, 2006.

#### Mahalanobis distance for multi-view pose estimation

We would like to use the Mahalanobis distance to measure reprojection errors of multi-view pose estimates; this distance can then provide a metric for the quality of pose estimates without ground truth labels.

This metric will be computed one body part at a time (across all camera views). Assume we have *V* camera views. For a given instant in time there will be an (*x, y*) prediction across all *V* views; stack these values into a single vector **x** = [*x*_1_, *y*_1_, …, *x*_*V*_, *y*_*V*_] ∈ ℝ^2*V*^. This 2*V* -dimensional vector represents a point in 3D space, so we can model it with the linear latent variable model of Eq. 4. The unique approach here is that instead of learning a single covariance matrix *D* for all observations, we will utilize observed ensemble variances that change from one observation to the next.

Now, if we consider the posterior predictive variance as defined in Eq. 8, the resulting *Q* would represent a single, joint measure of discrepancy across all camera views simultaneously. However, we would like to compute the posterior predictive variance *Q*^*v*^ for a single view *v* that incorporates information from the other views. Conditioning on the observations from the other views is straightforward in a linear model. Let us define *D*^*v*^ ∈ ℝ^2×2^ as the diagonal block of observed variances in *D* for view *v*; and *W*^*v*^ ∈ ℝ^2×3^ as the two rows of the loading matrix *W* that correspond to view *v*. Then

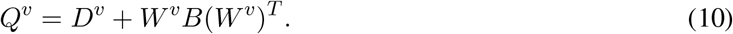

Finally, if we define **x**^*v*^ = [*x*_*v*_, *y*_*v*_] to be the observations for view *v*, and 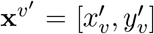 to be the expected observations for view *v* given observations from all remaining views (Eq. 8), then

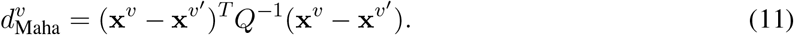

Concretely, the variance inflation procedure is carried out for each keypoint in each view at each point in time. For a given keypoint and timepoint, the value 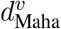 is computed, which measures the discrepancy between the observation in view *v* and the expected observation from all remaining views. This distance is on a standardized scale, the multi-dimensional equivalent of a *z*-score. If the distance is larger than 5, the prediction from view *v* is considered an outlier, and the variance is multiplied by a factor of 10 to reduce its influence on the other views in EKS. 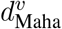 is recomputed, and this process of variance inflation continues until 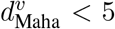. The final effect is inflating the variance in a view-specific block of the covariance matrix *D*_*t*_ in Eqs. 2 or 3.

#### Special case: two camera views

The above is a general procedure that becomes more robust as the number of camera views *V* increases. However, with only two views (*V* = 2), we face a potential indeterminacy issue. When predictions from both views are inconsistent yet each has low ensemble variance, it is impossible to determine which view (if either) is correct. Therefore, in the two-view case, if the Mahalanobis distance exceeds our threshold for one view, we inflate the variance in both views rather than trying to identify the problematic view.

### 4.11 Pseudo-label and distillation pipeline

The distillation pipeline converts mvEKS predictions (coordinates and uncertainty estimates) into high-quality pseudo-labels for retraining a single efficient model. Frame selection proceeds in two stages designed to ensure both *prediction quality* and *pose diversity* (Extended Data Fig. 6a). The pipeline is compatible with any multi-view pose estimator and operates regardless of whether camera calibration is available or not.

#### Stage 0: Inference and post-processing

We train ensembles of three networks using different train/validation frame splits. Larger ensembles will lead to better performance (Biderman et al., 2024). Once training is complete, inference is run on videos from the training set for each network. Predictions from all ensemble members are then post-processed with mvEKS to obtain 2D keypoint coordinates with uncertainty estimates.

#### Stage 1: Quality-based filtering

Frames are first filtered based on the mvEKS posterior predictive variance (*Q* in Eq. 9), which provides a measure of prediction uncertainty across all keypoints and views. To quantify instance-level confidence, we compute the maximum variance 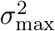 across all keypoints and views. Frames are ranked by 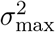, and the *N*_*f*_ frames with the lowest values are retained, prioritizing predictions with the lowest variance. We set *N*_*f*_ to ∼60% of the total frames in each video.

#### Stage 2: Diversity-based filtering

The subset of high-confidence frames is then subjected to clustering in 3D pose space to promote diversity. When camera parameters are available, 3D poses are obtained via triangulation; otherwise, PCA-based projection is used. *k*-means clustering is performed to identify representative poses (we use *k* = 3000), and for each cluster *j*, we select the frame closest to its center. The final selected frames are then converted into pseudo-labels using the original 2D mvEKS predictions, which are then used to retrain a single efficient model.

### 4.12 IBL: unsupervised clustering

Pose estimation was performed on the IBL mouse dataset using two model configurations compared in this work: (1) the ensemble median of three ResNet-50 networks, followed by triangulation; (2) and three LP3D networks post-processed with linear mvEKS (LP3D+EKS). All networks were trained with 200 labeled frames. The triangulation/EKS steps yield 3D trajectories for two keypoints: the left forepaw (`pawL`) and the right forepaw (`pawR`). All analyses were carried out across *n* = 39 sessions, each comprising 60,000 frames.

#### Kinematic feature extraction

For each time point in a session, an 11-dimensional kinematic feature vector was computed from the 3D trajectories of `pawL`and `pawR`. The feature vector comprised: (1) the 3D velocity vectors of each paw ((*v*_*x*_, *v*_*y*_, *v*_*z*_) for each; 6 dimensions), capturing both the direction and magnitude of paw movement; (2) the scalar speed of each paw (Euclidean norm of the velocity; 2 dimensions); (3) the scalar acceleration magnitude of each paw (Euclidean norm of the second-order finite difference; 2 dimensions); and (4) the inter-paw Euclidean distance (1 dimension), encoding instantaneous postural configuration. This yields a feature matrix of shape (*N*_frames_ × 11) per session per model.

#### Feature pooling and normalization

Feature matrices were computed independently for each session and model, then pooled into a single dataset. This allowed us to analyze behavioral states shared across models. Frames containing `NaN`values were excluded. The pooled matrix was standardized using robust scaling (median centering and interquartile range normalization), which is resistant to outliers.

#### Hierarchical agglomerative clustering

Behavioral states were identified by applying BIRCH (Balanced Iterative Reducing and Clustering using Hierarchies; Zhang et al., 1996) clustering to the full pooled, scaled feature matrix. The number of clusters was set to *k* = 10. To visualize the hierarchical relationships among the discovered behavioral states, a post-hoc linkage matrix was computed using Ward linkage on the cluster centroids; the resulting dendrogram illustrates the hierarchical relationships among the cluster centroids.

Each cluster was characterized by computing a robust *z*-score for each kinematic feature. Let **x**_*k,f*_ denote the set of values of feature *f* across all frames assigned to cluster *k*. The cluster deviation score is defined as:

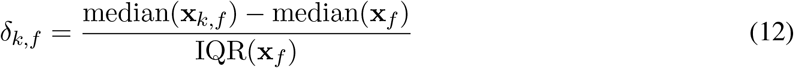

where median(**x**_*f*_) and IQR(**x**_*f*_) are the median and interquartile range of feature *f* computed over all frames in the dataset. This normalization is robust to the severe class imbalance present in behavioral data, where the large majority of frames correspond to quiescent states. Clusters were assigned interpretable labels based on the pattern of *δ*_*k,f*_ values across features (e.g., slow move, fast move, etc.).

#### Low dimensional embedding

To visualize the structure of the behavioral feature space, low-dimensional embeddings were computed using UMAP (McInnes et al., 2018) on a randomly drawn subsample of frames from the scaled 11-dimensional feature matrix. UMAP was run with `n_neighbors`= 199, `min_dist`= 0.3, and 2 output components, with a fixed random seed for reproducibility. Embedded points were colored by cluster assignment to verify that the discovered clusters occupied coherent, well-separated regions of the feature manifold.

### 4.13 IBL: neural decoding

To evaluate the downstream utility of each pose estimation model, we performed linear neural decoding of trial-based 3D paw kinematics from multi-region Neuropixels (Jun et al., 2017) recordings across *n* = 39 sessions from the IBL dataset.

#### Trial filtering

A trial is considered “valid” if it has no missing values in key event timestamps (stimulus onset, choice, feedback, first movement), the mouse made a choice, and reaction time falls within 0–10 seconds. Trials are further filtered to ensure all required behavioral signals (wheel speed and pose estimation) could be successfully extracted for that trial’s interval. Sessions contained between 127 and 390 valid trials.

#### Neural data processing

Single-unit spiking activity was binned into 20 ms bins over a 2 s window starting 500 ms before stimulus onset (`stimOn_times`), yielding a per-session tensor of shape (*n_trials* × 100 bins × *n_neurons*). Neurons from all recorded brain regions and all probes were pooled, and we did not filter neurons based on firing rate or other quality control metrics. Sessions contained a median of 1189 neurons, with a minimum of 274 neurons and a maximum of 2521; these neurons spanned a median of 10 Beryl-level brain regions per session, with a minimum of 4 regions and a maximum of 19 regions. The 39 sessions covered a collective 122 unique brain regions. Spike counts were *z*-scored across trials independently for each neuron and time bin, using mean and standard deviation estimated from the training partition only.

#### Behavioral target: 3D paw speed

We decoded the 3D speed of the left and right paws separately. For each session, 2D pose predictions were generated for each paw from three different models: (1) a ResNet-50 backbone trained on single-view data with median ensemble aggregation (ResNet-50); (2) LP3D with median ensemble aggregation (LP3D); and (3) an ensemble of three LP3D models post-processed with mvEKS (LP3D+EKS). Camera calibration parameters were used to triangulate the 2D predictions from both views into 3D world coordinates. Speed values were assigned to the temporal midpoint between consecutive frames and were retained in their original units without further normalisation. The resulting per-frame speed timeseries was then linearly interpolated to align with the neural trial windows.

#### Decoding model

We used ridge regression (using `sklearn`‘s `Ridge`class) to map neural population activity to the continuous paw speed timeseries. For each trial, the spike count matrix (100 bins × n_neurons) was flattened into a single vector and used to predict the 100 aligned target values. The regularisation strength *α* was selected via nested cross-validation using a grid search over *α* ∈ {10^−4^, 10^−3^, …, 10^4^}, with *R*^2^ as the scoring criterion.

#### Cross-validation

A strict 5-fold cross-validation scheme was applied such that each trial appeared in the held-out test set exactly once. In each fold, 80% of trials were used for training (with nested hyperparameter selection), and 20% formed the held-out test set. Within the training partition, data were further split 87.5%/12.5% into train and validation subsets for hyperparameter search. The final model was retrained on the full 80% training partition using the selected *α* before evaluation on the held-out test fold.

#### Differences with decoding analysis in the IBL Brainwide Map paper

Our decoding analysis differs from the Brainwide Map (BWM; IBL et al., 2025a) in several ways, reflecting differences in scientific goals and practical constraints.

First, to facilitate running multiple pose estimation and decoding models at scale, we restricted our analysis to 39 sessions (the BWM contains 459 sessions) and used only the first 100,000 frames (∼ 28 minutes) of each session, resulting in fewer valid trials per session (127–390 in our study; 128–1414 in the BWM).

Second, we relaxed the reaction time filter to include more trials, accepting reaction times between 0–10 s compared to the 0.08–2.0 s window used in the BWM.

Third, we used a slightly different and longer window of neural activity and behavior: a 2 s window starting 500 ms before stimulus onset time, rather than a 1.2 s window starting 200 ms before first movement time, following previous video-based decoding efforts with the BWM data (Zhang et al., 2026).

Fourth, we pooled all neurons within each session rather than splitting by brain region or filtering for quality. The BWM split neurons by region because that study aimed to characterize how different brain areas encode different task variables, a question we are not addressing here. The BWM also restricted to neurons passing a set of quality control metrics because it needed to compare results across analysis types (e.g., encoding and decoding) that imposed stricter requirements; we opted to use all available neural information since our goal is to compare the quality of different pose estimation models as decoding targets.

Fifth, the BWM decoded wheel speed and velocity rather than video-derived behavioral traces such as paw speed. This choice was partly due to missing or corrupted video data in some BWM sessions, and partly because—as evidenced in this manuscript—video-based paw tracking was not yet of sufficient quality for reliable downstream analysis.

Sixth, we do not perform the rigorous null-distribution-based statistical testing used in the BWM, which required permuting trials hundreds of times per session and target to construct null distributions. That procedure was necessary because the BWM asked whether a particular brain region significantly decodes a given target variable; our question is instead comparative: which of two pose estimation models produces a better estimate of the same intended target variable (paw speed). We therefore report direct *R*^2^ values (Table 2, Fig. 4) rather than the null-adjusted *R*^2^ values used in the BWM.

## 5 Data availability

We will make all labeled data publicly available upon manuscript acceptance.

## 6 Code availability

The Lightning Pose code is available at https://github.com/paninski-lab/lightning-pose under the MIT license. Run pip install lightning-pose to install the latest release of Lightning Pose via the Python Package Index (PyPI).

The Ensemble Kalman Smoother code is available at https://github.com/paninski-lab/eks under the MIT license. Run pip install ensemble-kalman-filter to install the latest release.

The user interface code is available at https://github.com/paninski-lab/lightning-pose-app under the MIT license. This code enables launching our app locally, or on cloud resources by creating a Lightning.ai account. Run pip install lightning-pose-app to install the latest release.

## 7 Extended Data Figures

**Extended Data Figure 1:**
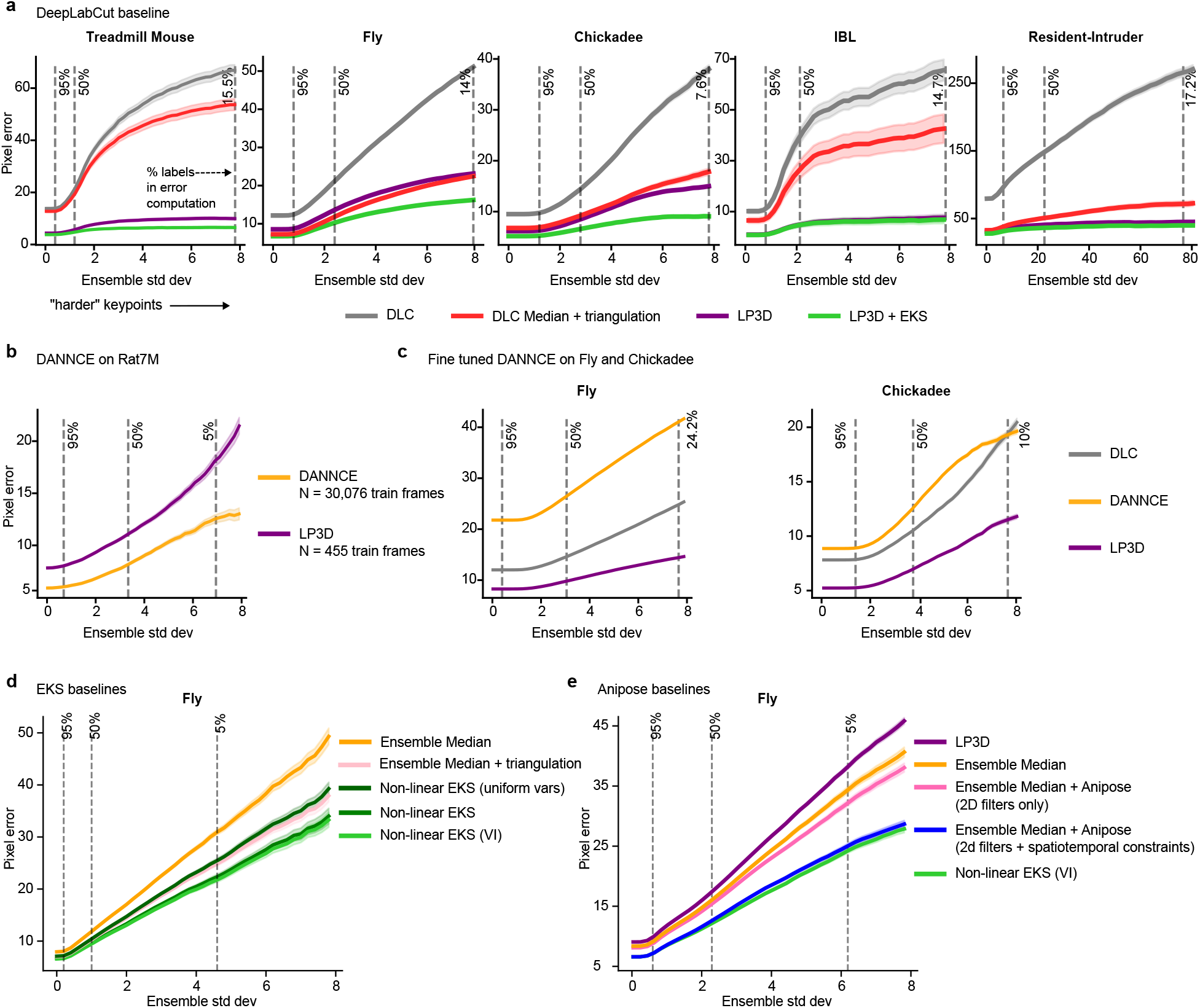
LP3D outperforms baselines across datasets. **a**, Pixel error for all models trained on 200 labeled frames, three seeds per model. Individual DeepLabCut (DLC; Mathis et al., 2018) networks (gray) represent the standard single-view baseline. The DLC ensemble combined with post-hoc triangulation (red) improves upon individual DLC networks by aggregating predictions across views (for Treadmill Mouse, where camera parameters are unavailable, we use 3D PCA in place of triangulation). LP3D (purple) substantially outperforms individual DLC networks and performs comparably to or better than DLC + triangulation, demonstrating that joint multi-view processing at the representation level is competitive with post-hoc geometric fusion. LP3D + EKS (light green) achieves the best performance across all datasets, demonstrating the additional benefit of uncertainty-aware post-processing that combines both temporal smoothing and geometric consistency (for Treadmill Mouse we apply the linear EKS variant; all other datasets use the nonlinear variant). **b**, DANNCE (Dunn et al., 2021; orange), pretrained on 30,076 Rat7M frames, slightly outperforms LP3D trained on only 455 labeled frames across 96 held-out test frames. **c**, DANNCE fine-tuned on the Fly and Chickadee datasets (three seeds, 200 labeled frames each). LP3D significantly outperforms DANNCE on both datasets, as does DLC. **d**, Comparison of EKS variants on the Fly dataset. Ensemble median + triangulation (light pink) performs comparably to nonlinear EKS with uniform variances (dark green), indicating that per-frame and -keypoint ensemble variances contribute to improved performance. Nonlinear EKS with variance inflation (light green) achieves the best results. **e**, Nonlinear EKS with variance inflation performs similarly to the full Anipose pipeline on the Fly dataset (Karashchuk et al., 2021; see Sec. 4.7 for details).

**Extended Data Figure 2:**
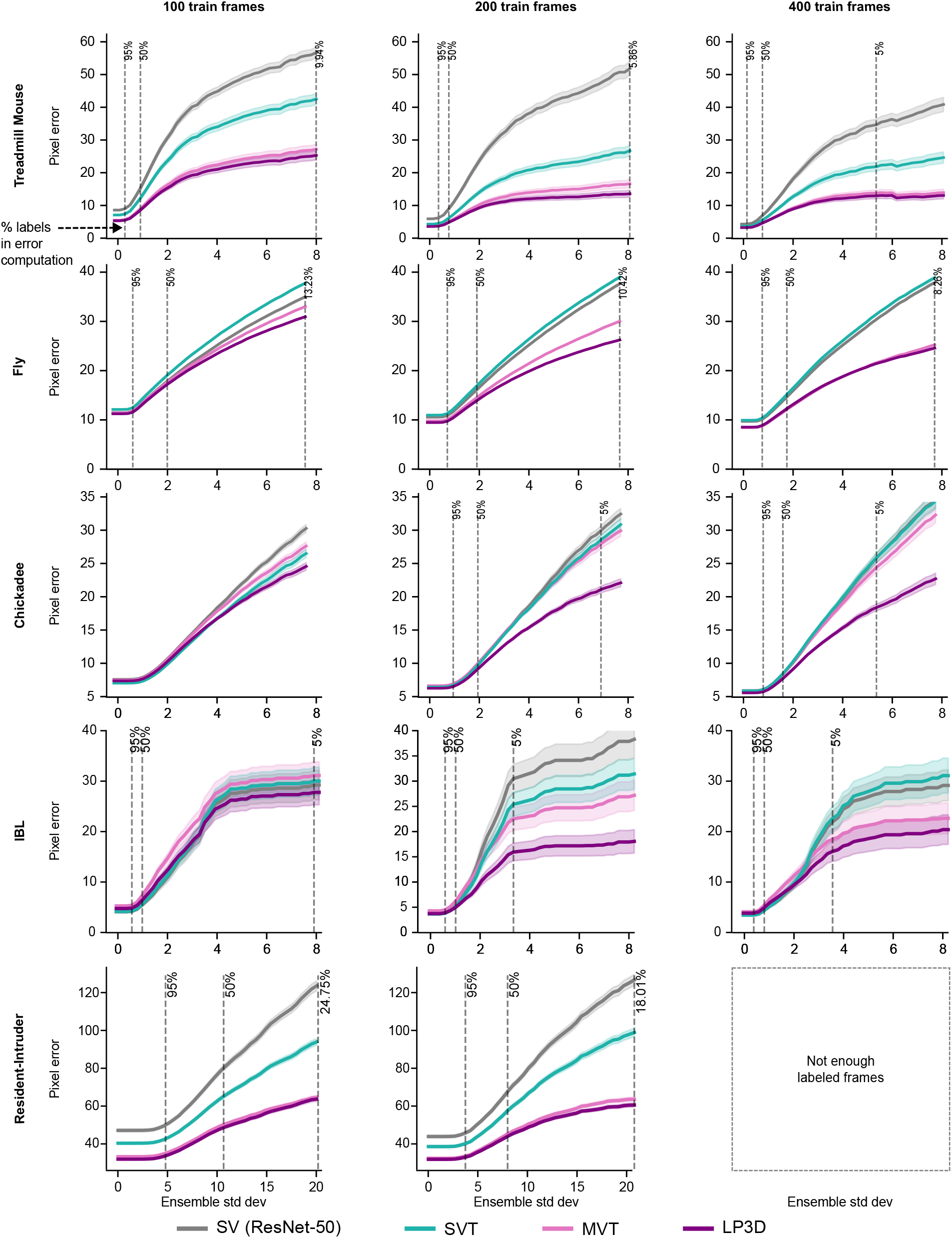
Model performance as a function of labeled data. An ensemble of three networks with different train/validation splits are trained for each model type. Rightmost results for Fly use the maximum of 377 labeled frames. Resident-Intruder only has 207 labeled training frames.

**Extended Data Figure 3:**
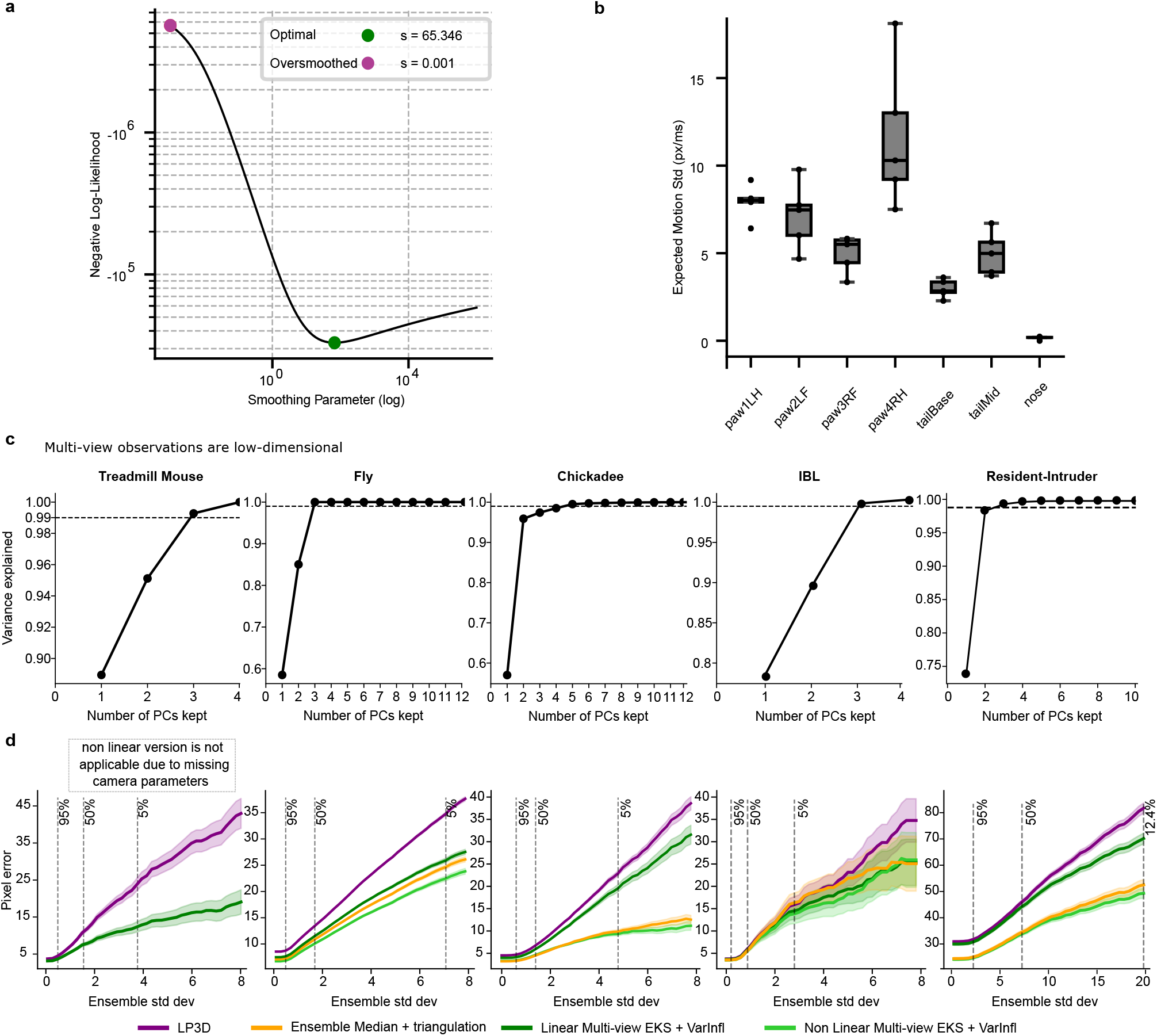
Nonlinear Ensemble Kalman Smoother improves pose estimation. **a**, Log-likelihood as a function of mvEKS smoothing parameter shows a single well-defined optimum. **b**, Expected motion magnitude varies by keypoint in the Treadmill Mouse dataset: the nose exhibits low expected motion (high temporal consistency), while the paws show greater expected variability in position across frames (pixels per millisecond). **c**, Multi-view observations are low-dimensional. Variance explained for increasing numbers of Principal Component dimensions for each dataset. Three dimensions explain more than 99% of the variance for the Treadmill Mouse, Fly, IBL, and Resident-Intruder datasets, while the Chickadee dataset (which includes more camera distortion) requires five dimensions to exceed 99% variance explained. **d**, Calibrated post-processor comparison. Variance-inflated nonlinear mvEKS (light green) outperforms both variance-inflated linear mvEKS (dark green) and simple triangulation run on the ensemble median (i.e., same input as the mvEKS models; orange). Nonlinear mvEKS is not applicable to the Treadmill Mouse as it lacks camera parameters. However, we can still run the linear version of mvEKS (dark green) on this dataset, which is not possible with standard triangulation based post-processors like Anipose.

**Extended Data Figure 4:**
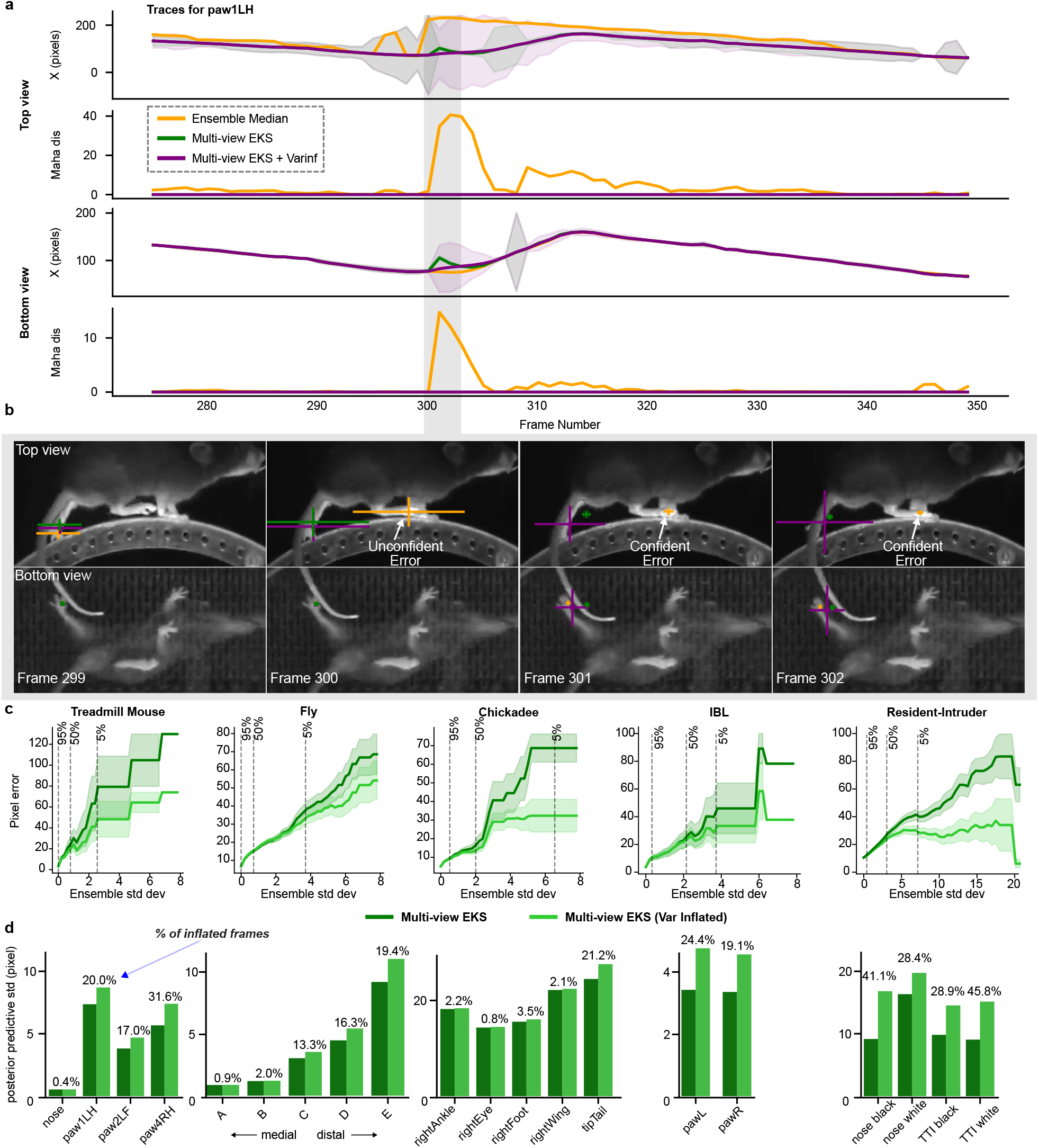
Variance inflation resolves cross-view inconsistencies. **a**, Example traces for one keypoint (Treadmill Mouse, left hind paw) from ensemble median, linear mvEKS, and linear mvEKS with variance inflation, top and bottom views. Error bands show posterior predictive variance from EKS. High Mahalanobis distance marks low confidence or cross-view disagreement. Occluded segment shaded in gray. Variance inflation (purple) improves temporal smoothness and increases uncertainty estimates. **b**, Frames 299–302 (occlusion in (a)). Ensemble median disagrees across views; mvEKS improves consistency (green; frames 300-302); variance-inflated mvEKS aligns both views and properly signals uncertainty (purple; frames 300-302). **c**, Variance-inflated mvEKS (light green) achieves lower pixel error than mvEKS (green) across keypoint difficulty levels. **d**, Variance inflation occurs most frequently on distal keypoints such as paws and limbs, consistent with their higher variability and susceptibility to occlusion. The specific keypoints affected, however, depend on recording geometry and behavioral context: in the IBL dataset, paw keypoints are difficult to resolve from both views simultaneously; in the Resident-Intruder dataset, nose keypoints and tail tip (TTI) are among the most frequently inflated, reflecting mutual occlusion during close social interactions. Percentage of keypoints with inflated variance annotated above each bar.

**Extended Data Figure 5:**
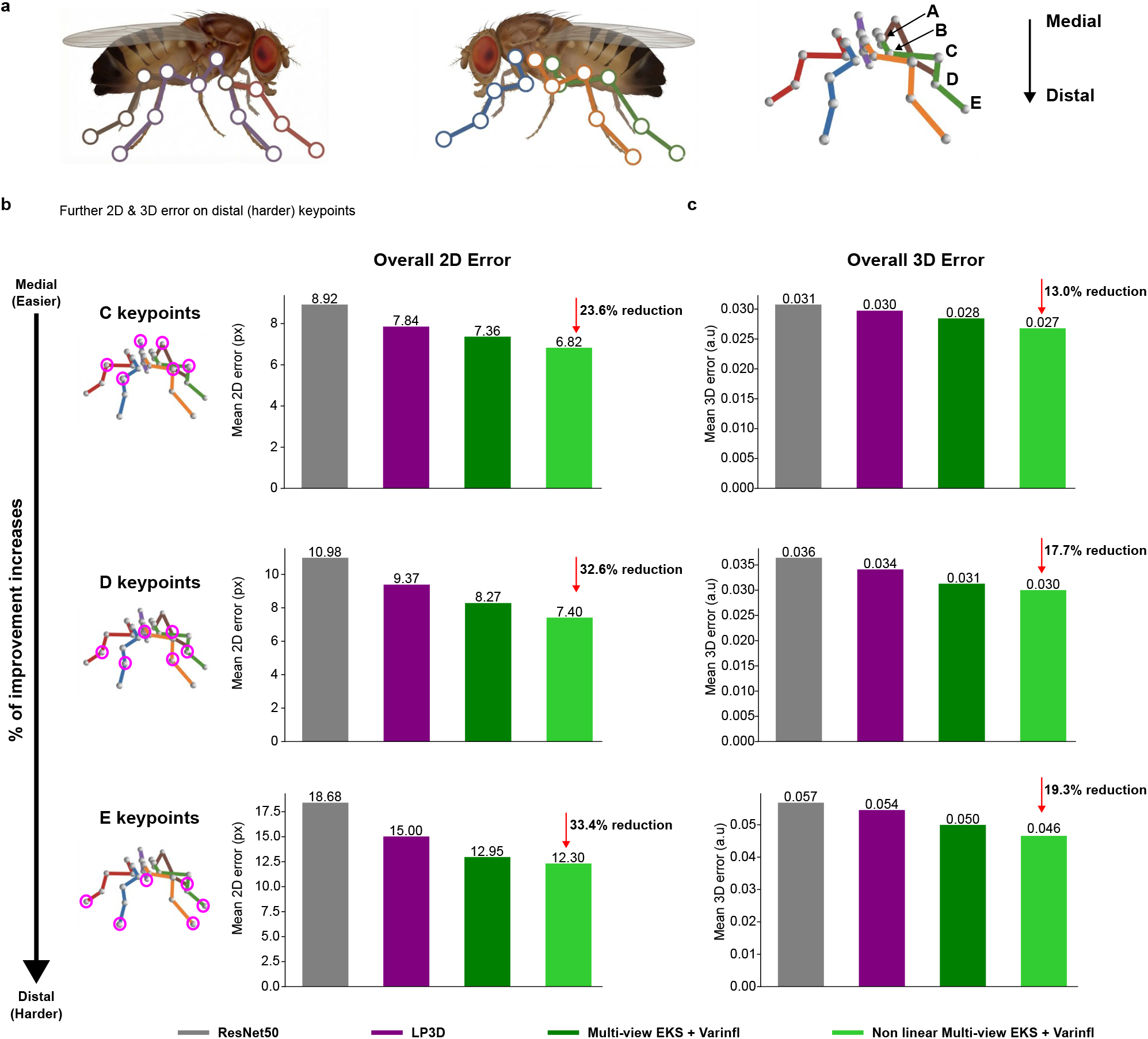
LP3D and EKS reduce error most for hardest Fly body parts. **a**, Fly behavior was filmed with six cameras evenly distributed around the floating ball. Five keypoints per leg (one per major joint; A–E from medial to distal) were labeled and tracked, giving six keypoints per joint type (one per leg). Distal keypoints are more challenging due to smaller size and more variable motion. **b**, 2D error from medial to distal (C–E) on held-out test frames. Error is averaged across views and across the six keypoints per joint type (one per leg). LP3D outperforms ResNet-50 across keypoints, with larger gains on the most challenging (distal) ones. Variance-inflated nonlinear mvEKS outperforms linear mvEKS. Red indicates percentage reduction in error relative to the ResNet-50 baseline. **c**, 3D error (averaged across keypoints) shows the same trend: variance-inflated nonlinear mvEKS achieves the best performance and has the largest gains on the most challenging body parts.

**Extended Data Figure 6:**
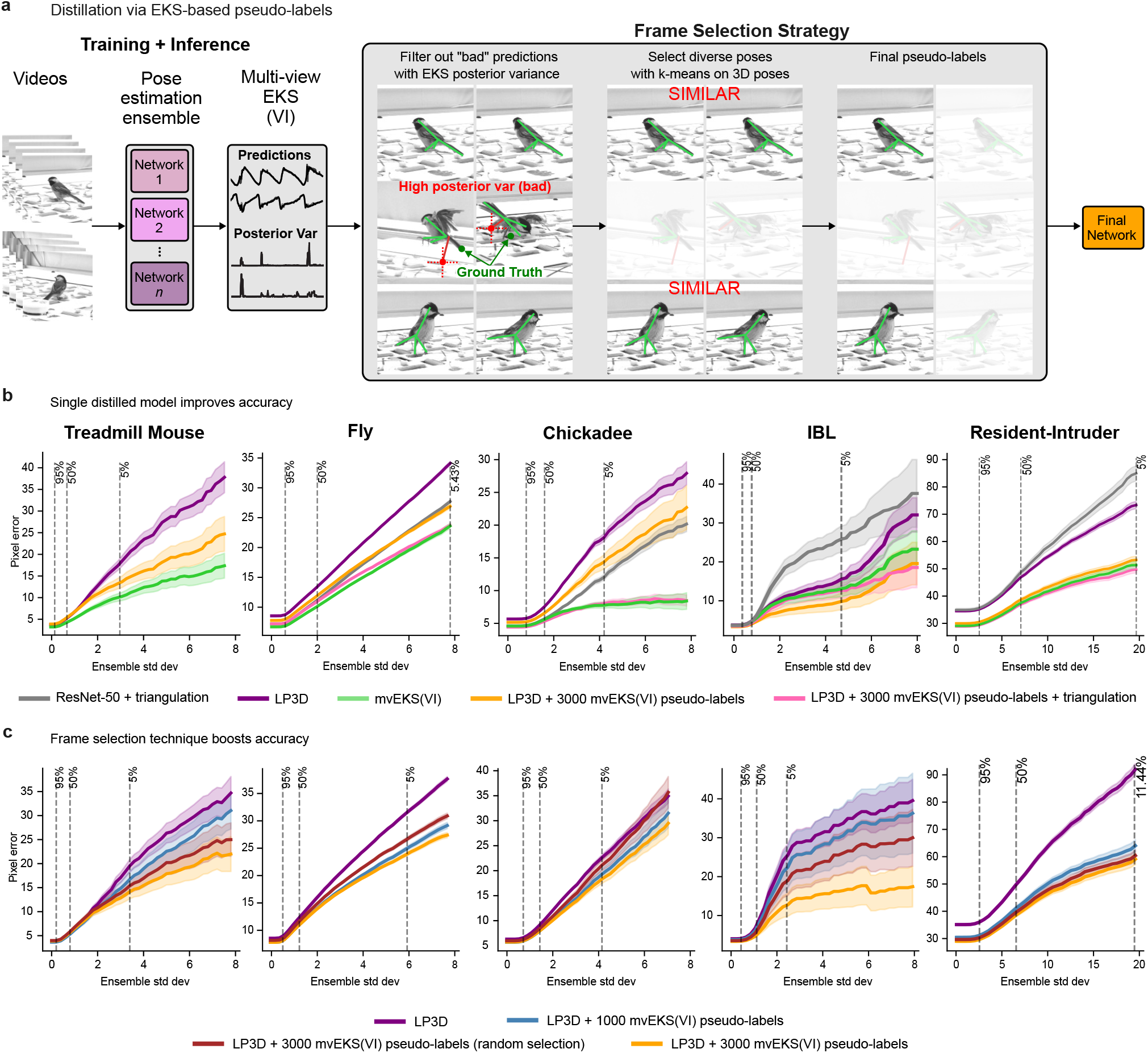
EKS-based pseudo-labels improve pose estimation. **a**, Schematic of our pseudo-label selection and training procedure. **b**, The distilled LP3D+EKS model (orange) outperforms initial ensemble member LP3D models (purple), but does not reach the performance of nonlinear EKS (green; Treadmill mouse uses linear EKS). Enforcing geometric consistency on the distilled model output (pink) brings single-model performance levels equal to that of the full LP3D+EKS pipeline (green). For calibrated setups, we also compare against the standard ResNet-50+triangulation baseline (gray). **c**, Targeted frame selection (yellow) outperforms random selection (red). Additional high quality labeled frames lead to better results (blue versus yellow).

**Extended Data Figure 7:**
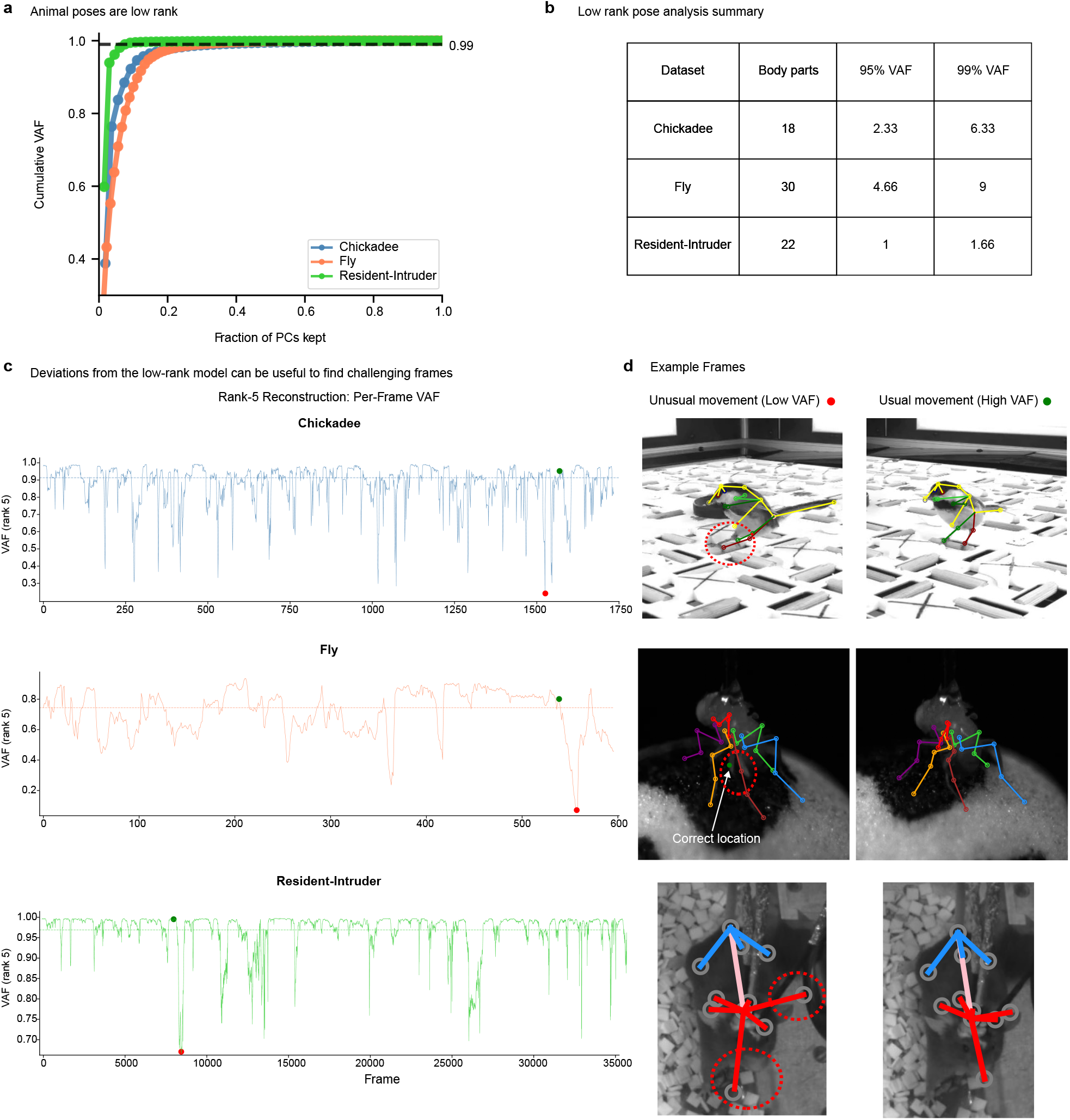
3D poses are low rank. **a**, Cumulative variance accounted for (VAF, %) versus fraction of PCs kept, computed on egocentrically-aligned 3D pose coordinates for Chickadee, Fly, and Resident-Intruder datasets. After egocentric alignment, poses for Chickadee and Resident-Intruder are low-rank; the Fly dataset is also low-rank without egocentric alignment. **b**, Summary of low-rank pose analysis. The number of components required to explain 95% and 99% of variance is normalized by the number of body parts (divided by 3, accounting for *x, y, z* dimensions), showing that fewer than 5 effective body parts explain 95% of pose variance. **c**, Deviations from the low-rank model can be useful to find challenging frames. Rank 5 reconstruction per-frame VAF shows challenging frames for all datasets; one representative challenging frame (red dot, *low* VAF) and one representative typical frame (green dot, *high* VAF) are shown for each dataset. **d**, Example frames corresponding to the red and green dots in (c). Frames with low rank-5 VAF (red) correspond to unusual or extreme poses that deviate from the dominant low-rank structure; frames with high VAF (green) correspond to typical, well-reconstructed poses. Red circles indicate incorrect predictions for the low VAF frames.

## 8 Supplementary Video Captions

**Supplementary Video 1**. Predictions from Ensemble Median, linear mvEKS, and linear mvEKS with variance inflation for the paw1LH and paw2LF keypoints, shown with error bars indicating uncertainty in the *x* and *y* directions, for the Treadmill Mouse dataset. Orange error bars indicate ensemble variance; green error bars indicate posterior variance from mvEKS. This session corresponds to the example shown in Extended Data Fig. 4a-b for paw1LH.

**Supplementary Video 2**. Baseline of ResNet-50 versus LP3D + mvEKS with variance inflation (200 labeled frames) for both paws, IBL dataset, similar to the example session in Fig. 4e. Red error bars indicate ensemble variance; green error bars indicate posterior variance from mvEKS. At frames 1070–1077 and frame 1084, LP3D + mvEKS outperforms ResNet-50, maintaining consistent tracking across views during occlusion events where ResNet-50 produces erroneous predictions. At frames 10188–10195, ResNet-50 outperforms LP3D + mvEKS, though both models produce reasonable predictions in this case.

**Supplementary Video 3**. LP3D (200 labeled frames) + mvEKS with variance inflation, Fly dataset. Error bars indicate posterior variance from mvEKS.

**Supplementary Video 4**. LP3D (200 labeled frames) + mvEKS with variance inflation, Chickadee dataset. Error bars as in Supp Video 3.

**Supplementary Video 5**. LP3D (200 labeled frames) + mvEKS with variance inflation, cropped on the black mouse from the Resident-Intruder dataset. Error bars as in Supp Video 3.

**Supplementary Video 6**. LP3D (200 labeled frames) + mvEKS with variance inflation, cropped on the white mouse from the Resident-Intruder dataset. Error bars as in Supp Video 3.

**Supplementary Video 7**. Comparison of DANNCE and LP3D model predictions on Fly dataset alongside ground truth labels (150 test frames) for 12 keypoints (2 medial joints per leg across all 6 legs), corresponding to the results shown in Extended Data Fig. 1c.

**Supplementary Video 8**. Comparison of DANNCE and LP3D model predictions on the Chickadee dataset alongside ground truth labels (143 test frames) for the left foot, right foot, top of head, and tail base key-points, corresponding to the results shown in Extended Data Fig. 1c.

## 9 Supplementary material

**Supplementary Figure 1:**
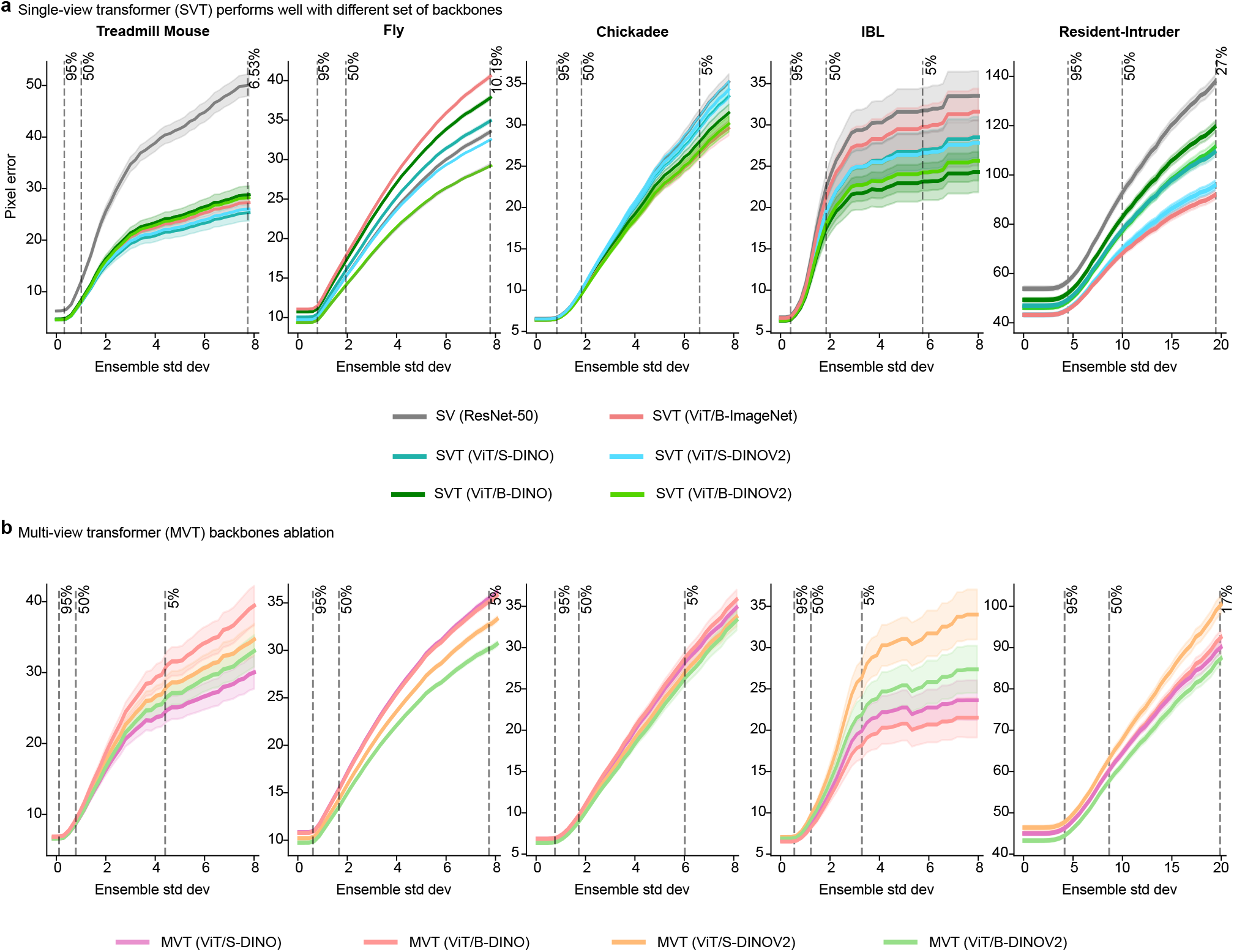
Comparison of pretrained transformer backbones and ResNet-50 for SVT and MVT. **a**, Comparing single-view transformer (SVT) performance with different transformer pretrained backbones by showing pixel error as a function of keypoint difficulty (lower is better). Dashed vertical lines indicate the percentage of data used for the pixel error computation. V_I_T/B is a “base” model (∼ 80M parameters), V_I_T/S is a “small” model (∼ 20M parameters); ResNet-50 has ∼20M parameters. ViT-B/ImageNet refers to ViT B-16 pretrained on ImageNet using masked autoencoders (MAE; He et al., 2022). DINO is a self-supervised pretraining method based on knowledge self-distillation with Vision Transformers (Caron et al., 2021); DINOv2 is an improved version trained on a larger curated dataset (Oquab et al., 2023). ViT-B/DINO and ViT-S/DINO denote base and small ViT models pretrained with DINO on ImageNet, respectively; ViT-B/DINOv2 and ViT-S/DINOv2 denote base and small ViT models pretrained with DINOv2. **b**, Comparing multi-view transformer (MVT) performance with different transformer pretrained backbones by showing pixel error as a function of keypoint difficulty.

**Supplementary Figure 2:**
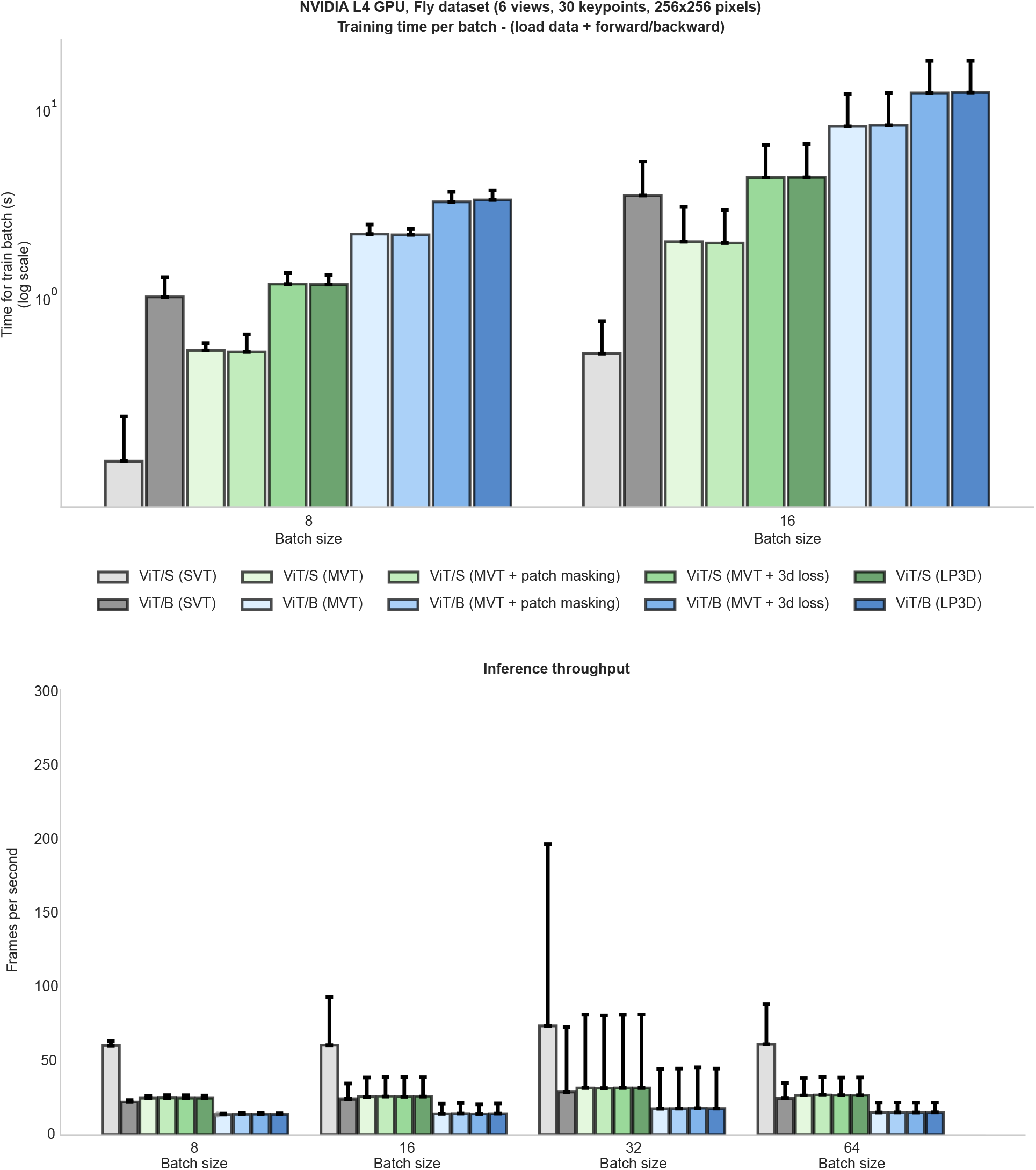
Training time per batch and inference throughput. Timing benchmarks are performed with the Fly dataset on an entry-level NVIDIA L4 GPU. Top panel: each bar depicts the mean batch processing time (in seconds) and 95% CI over *n* = 100 batches; *y*-axis is log-scaled. Bars are grouped by batch size (8/16) along the *x*-axis. Bottom panel: each bar depicts the mean frames per second with 95% CI over *n* = 100 batches. Bars are grouped by batch size (8/16/32/64). For both panels, 50 warm-up batches were processed before recording measurements.

**Supplementary Figure 3:**
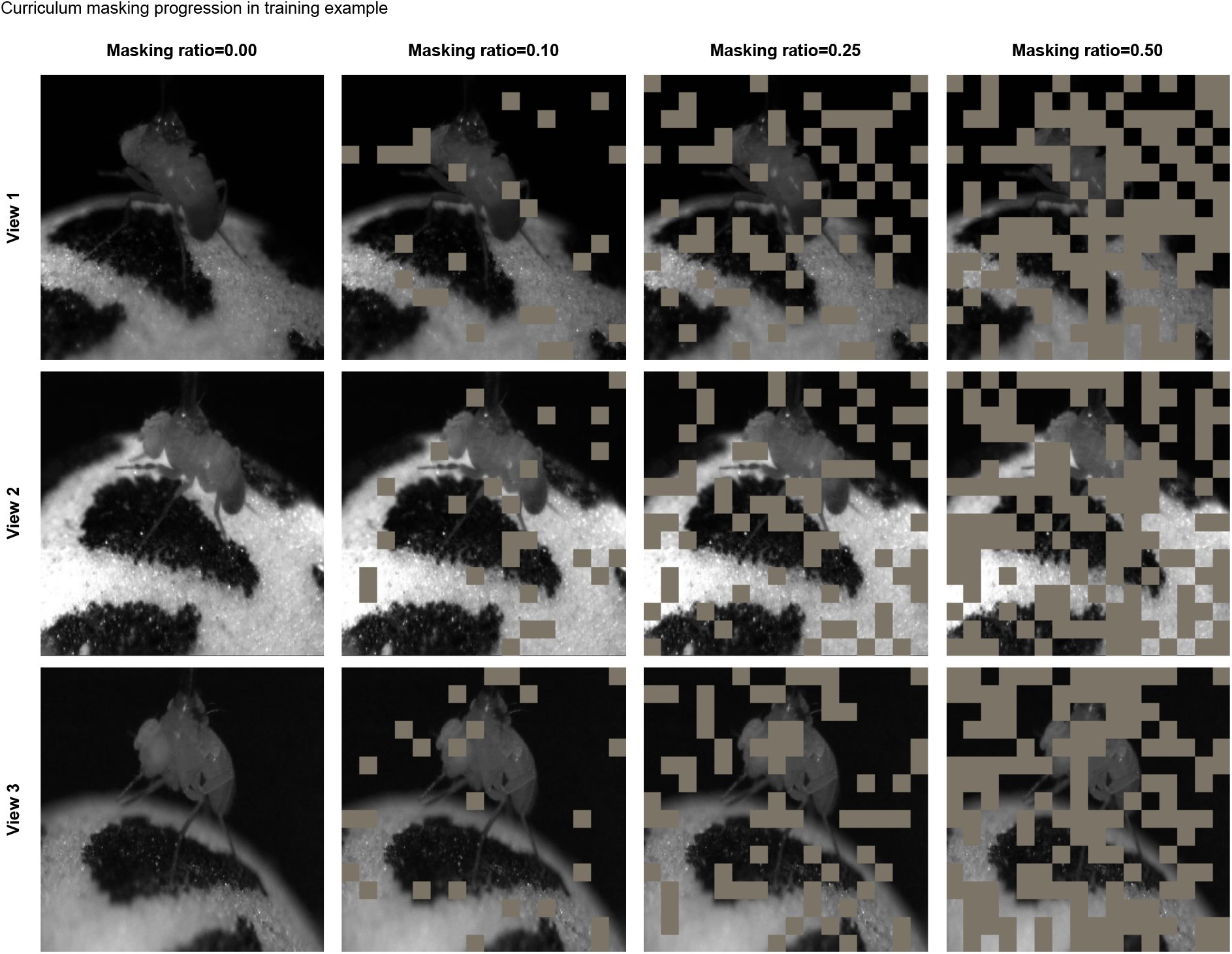
Simulating occlusions in multi-view transformer training via curriculum patch masking. Visualization of the patch masking scheme applied to three example camera views from the Fly dataset. Each 256 × 256 input image is divided into a regular grid of 16 × 16 pixel patches, and a fraction of patches are randomly zeroed out (brown regions). Columns show increasing masking ratios (0.00, 0.10, 0.25, 0.50) corresponding to the curriculum schedule used during training: the model first trains without masking, then the masking ratio is linearly increased from an initial to a final value over a defined range of training steps. By synthetically occluding portions of individual views, this scheme simulates the partial occlusions that frequently arise during natural animal behavior, forcing the model to leverage cross-view information through the self-attention mechanism of the multi-view transformer. The model thus learns to compensate for missing information in one view by drawing on unoccluded views, improving robustness to occlusions at inference time.

**Supplementary Figure 4:**
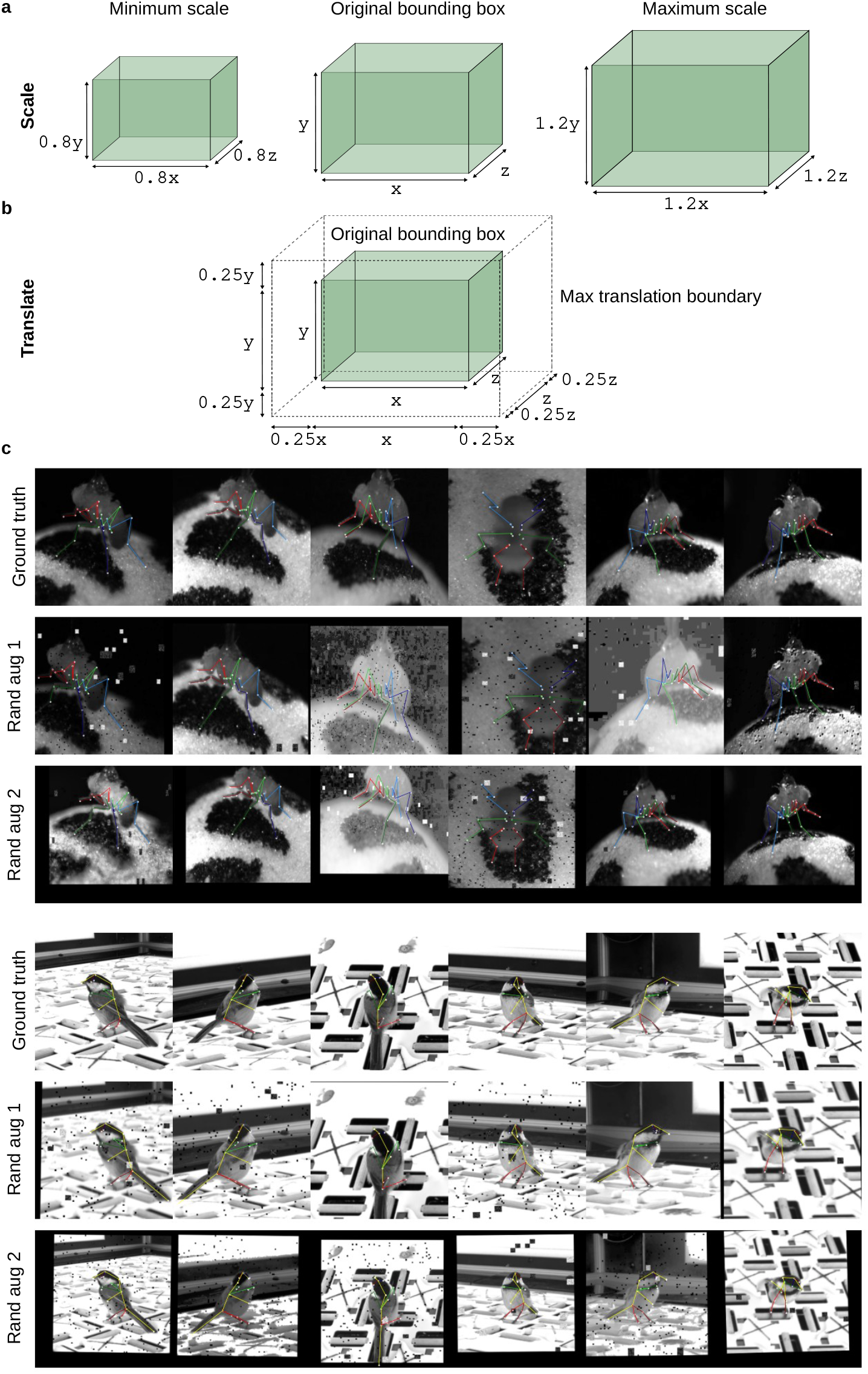
Illustration and examples of 3D augmentations. **a**, The *scale* augmentation samples a random scalar from [0.8, 1.2] and multiplies each centered keypoint by this value. This resizes the 3D bounding box while preserving the subject’s aspect ratio and maintaining the centered position. **b**, The *translate* augmentation samples random scalars from [− 0.25, 0.25] for each dimension, then multiplies these by the corresponding bounding box dimension length. This produces translation amounts that scale appropriately with subject size. **c**, Augmentations for datsets with camera calibration parameters combine scale and translation in the 3D space with view-independent appearance augmentations (e.g., pixel noise and brightness).

**Supplementary Figure 5:**
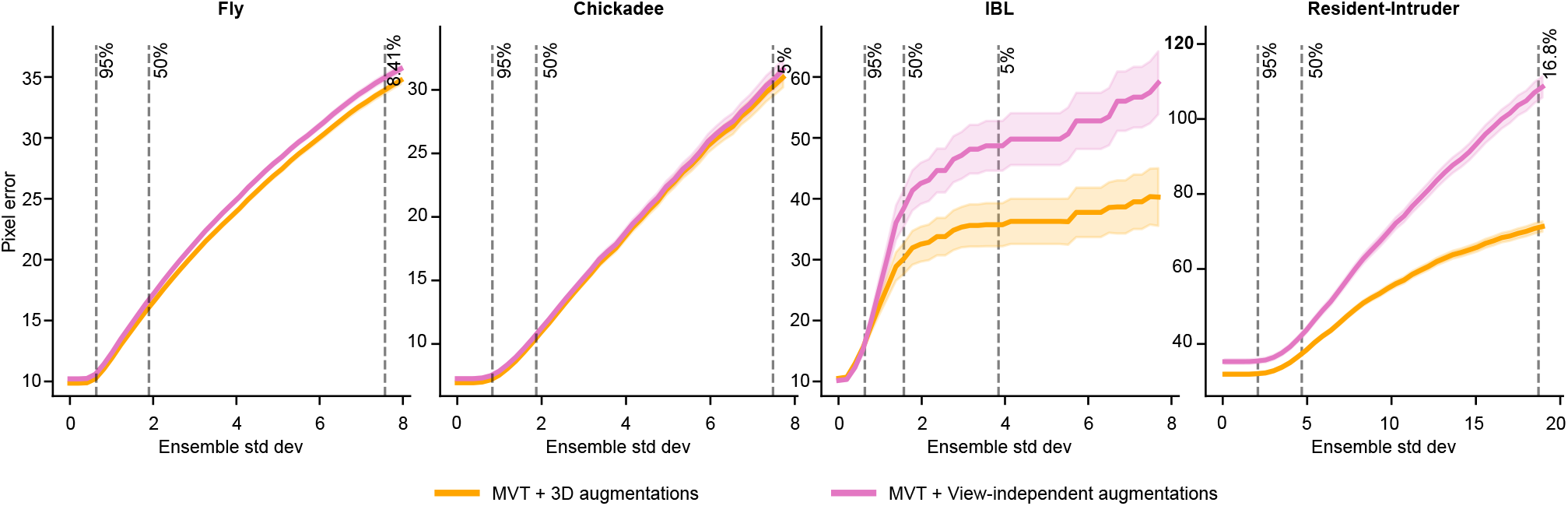
3D augmentations compare similarly or outperform view-independent augmentations. 3D augmentations, described in (Supp Fig. 4), can be applied to any dataset with camera calibration parameters. These geometrically consistent augmentations perform similar to or better than view-independent geometric augmentations, such as rotations and scales. Both forms of augmentation use the same appearance-based augmentation pipeline (pixel noise, brightness/contrast manipulations, etc.).

**Supplementary Figure 6:**
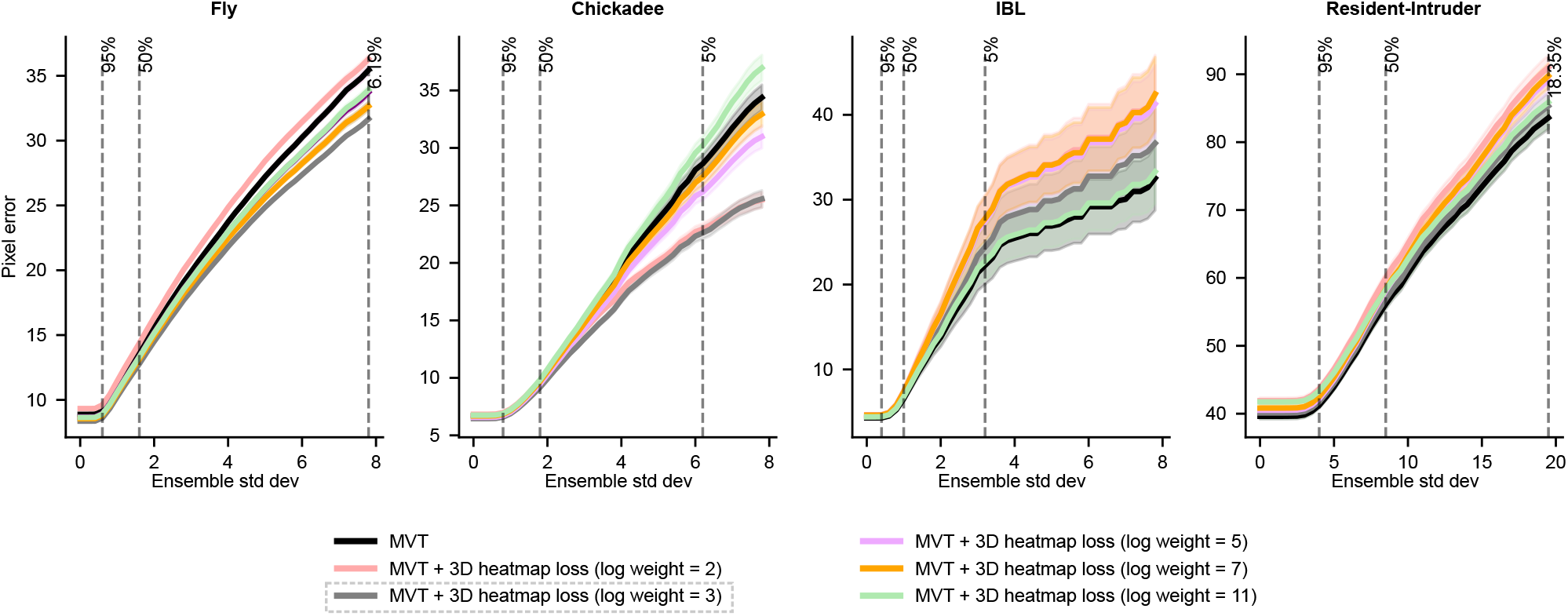
Hyperparameter selection for the 3D reprojection loss weight. Performance across datasets for different log weight values balancing the 3D reprojection loss against the 2D heatmap loss. Higher log weight values correspond to lower loss weights. The gray dashed line marks the selected value (`log weight=3`), which performs consistently well across all datasets and is used for all experiments.

**Supplementary Figure 7:**
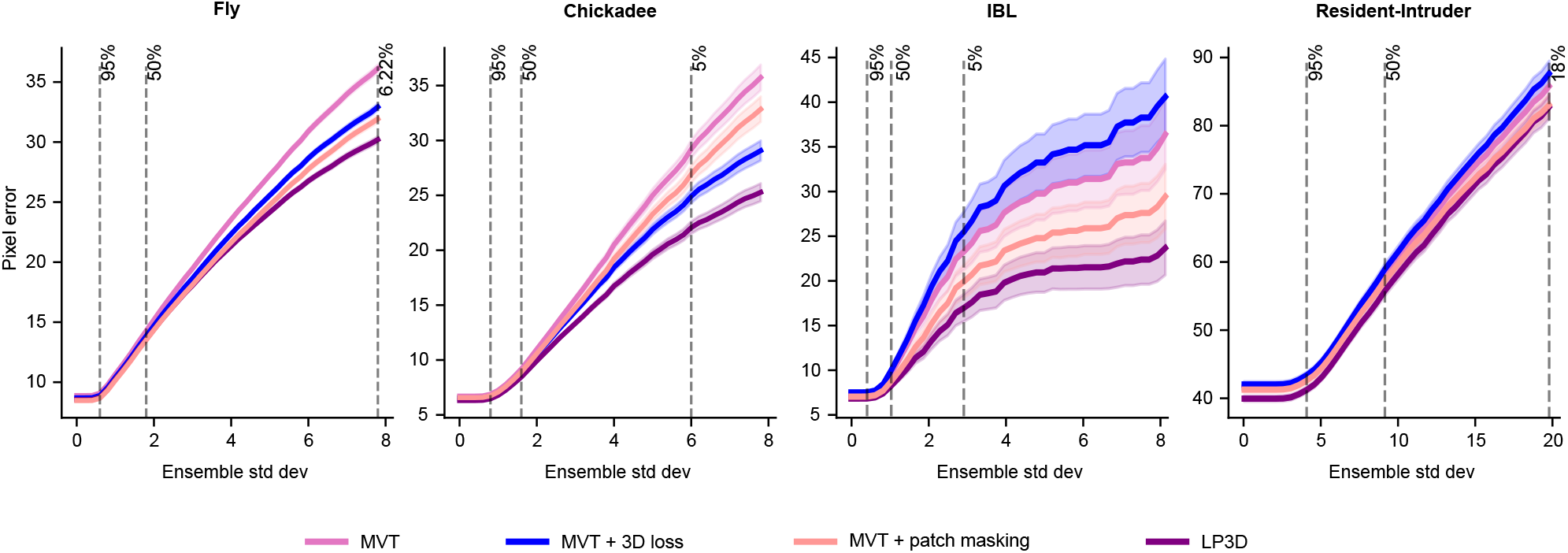
Patch masking and 3D reprojection loss provide complementary performance benefits. Performance across all datasets, comparing MVT alone; MVT with patch masking; MVT with the 3D reprojection loss; and the full model with both patch masking and 3D reprojection loss (LP3D). For Fly and Chickadee, each component independently improves over MVT alone. For IBL and Resident-Intruder, the 3D loss alone hurts performance relative to MVT, yet combining it with patch masking surpasses patch masking alone, demonstrating that the two components are complementary.

**Supplementary Figure 8:**
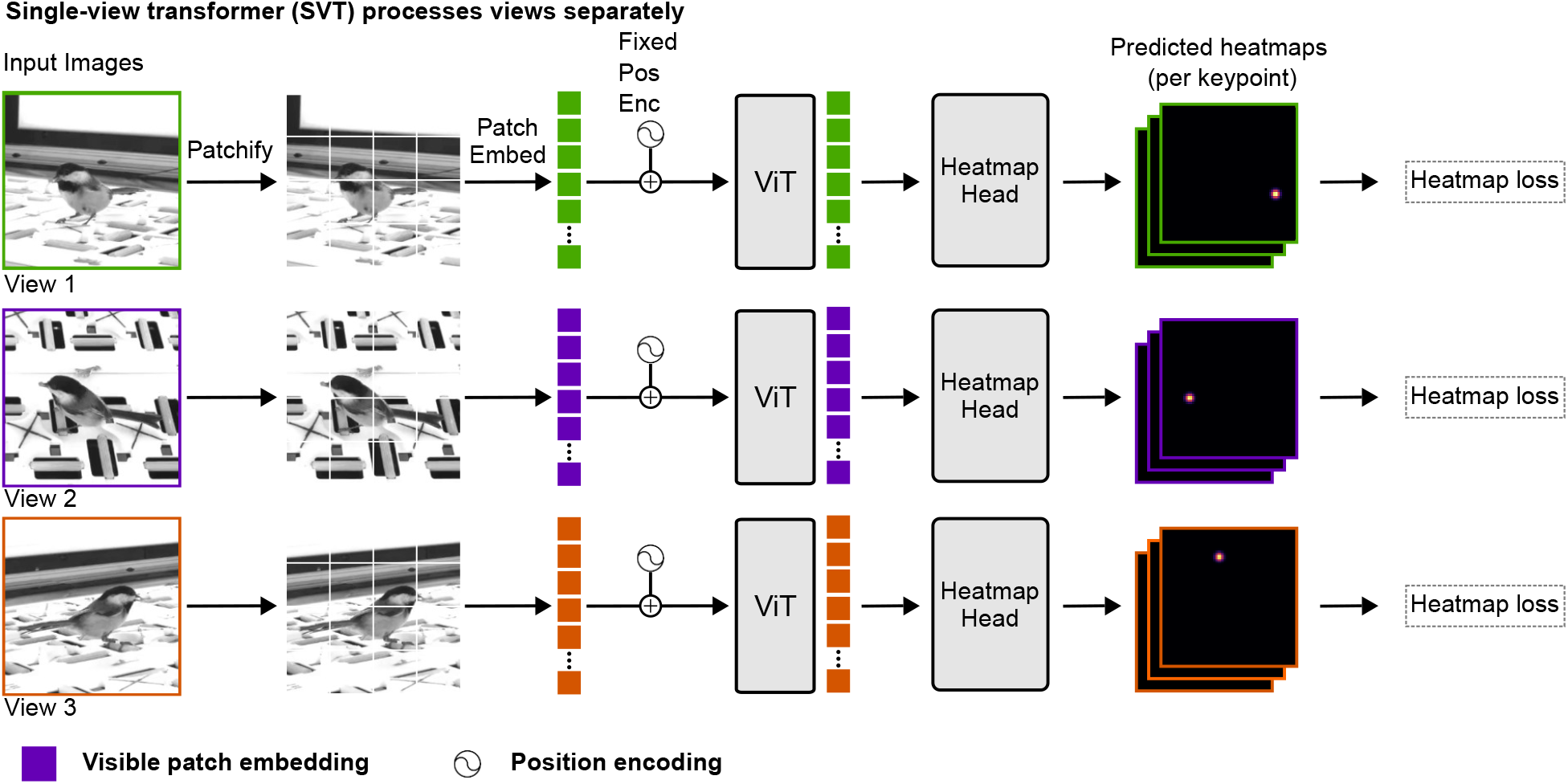
Single-view transformer (SVT) architecture. Input frames are split into patches, embedded into a latent space, combined with a fixed position encoding, and processed through a vision transformer (ViT). Outputs are reshaped and passed to a heatmap head. The model is trained with a mean square error (MSE) loss between predicted and ground truth heatmaps. Multiple views are processed independently.

**Supplementary Figure 9:**
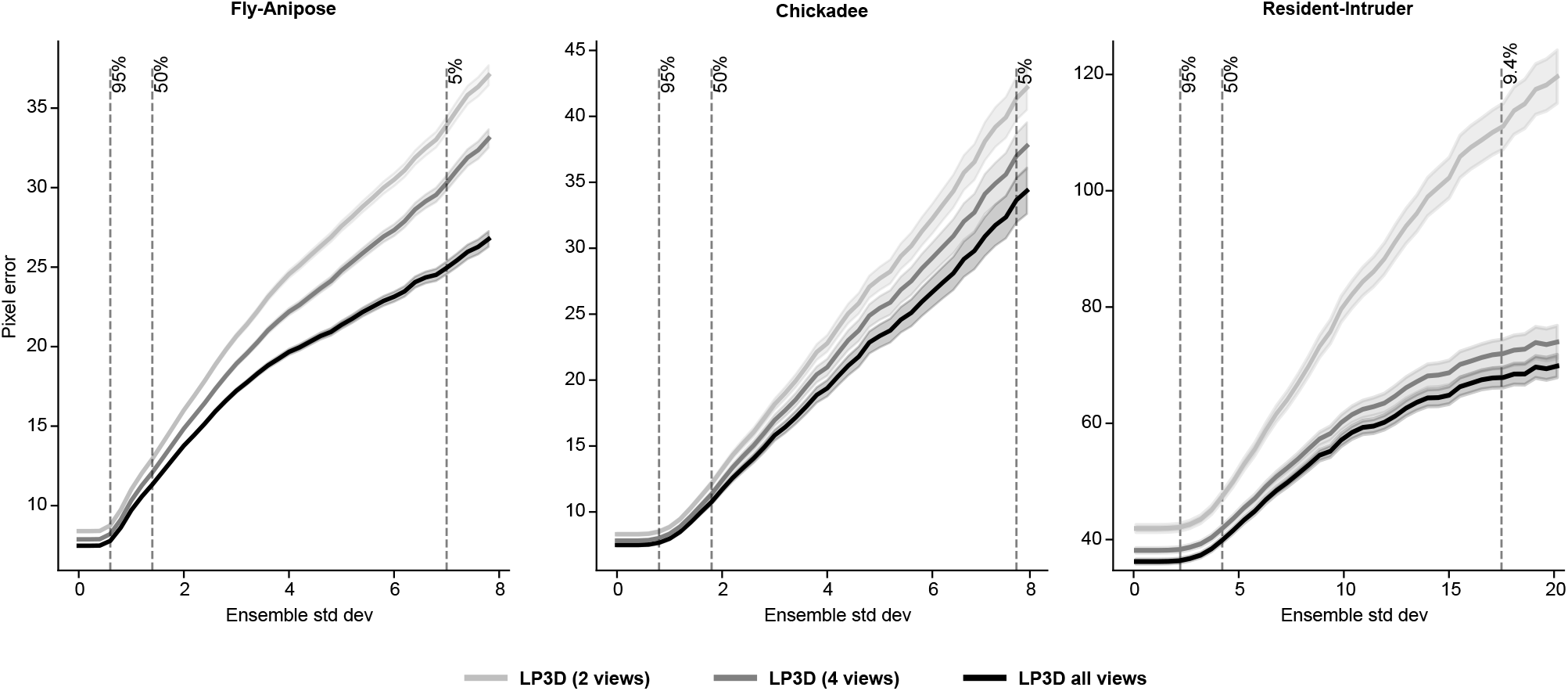
The accuracy of LP3D increases with the number of camera views. The LP3D model was trained on the Fly and Chickadee datasets (both six-view) and on the Resident-Intruder (five-view). We varied the number of views used during training to either two, four, or five/six views (three random seeds per condition). The two views used for the two-view condition were a random subset of the four views, which in turn were a random subset of all available views. All models were evaluated on the same shared two-view subset for 2D keypoint prediction (these two views are present in each view subset). This design demonstrates that increasing the number of views used during training significantly improves the LP3D model’s performance, as evaluation is held constant across all conditions.

**Supplementary Figure 10:**
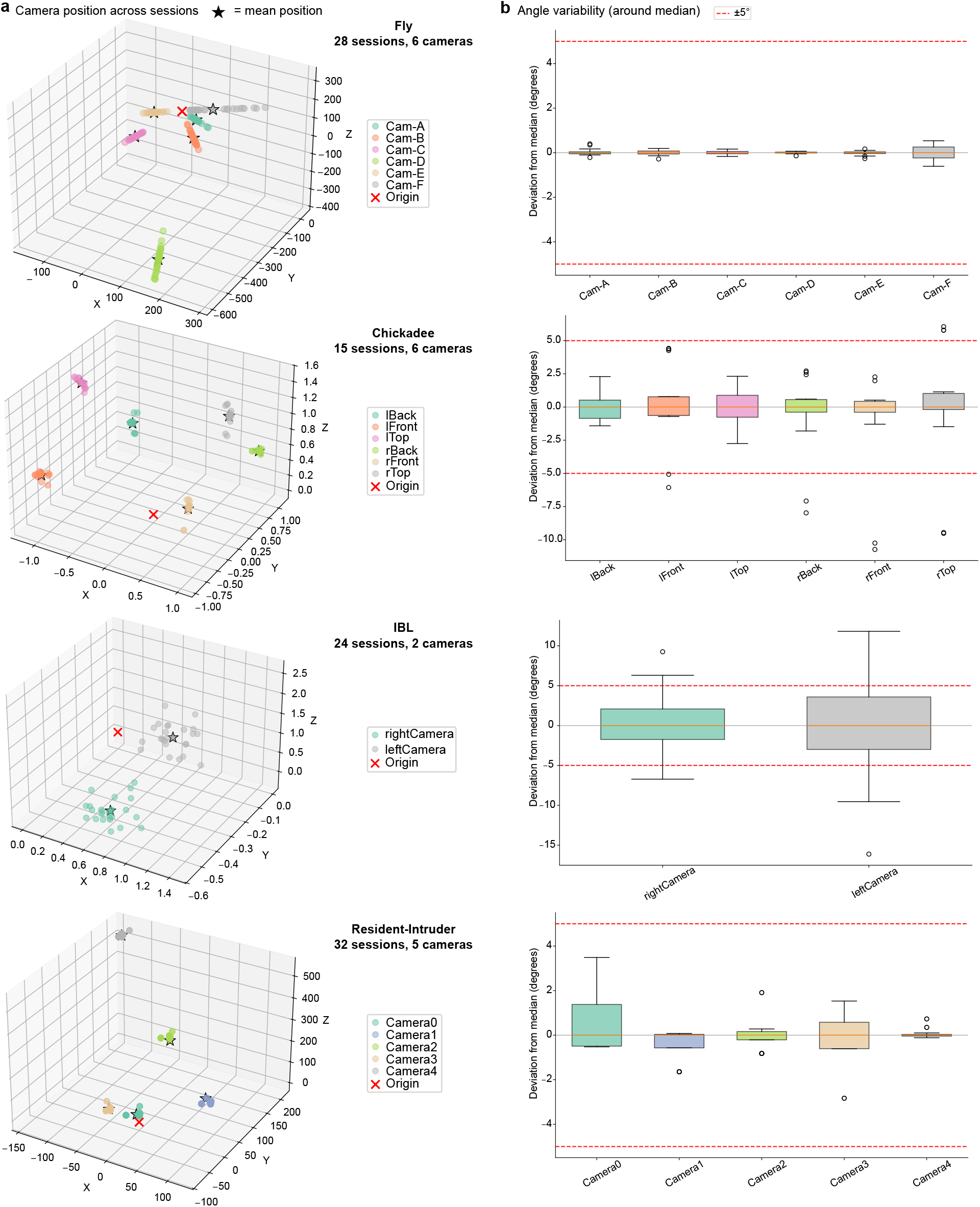
Camera stability across recording sessions. **a**, Three-dimensional positions of each camera’s optical center, estimated from per-session calibration files, for four benchmark datasets (Fly, Chickadee, IBL, Resident-Intruder). Each point represents one session; stars indicate the per-camera mean position. **b**, Deviation of each camera’s rotation angle (Rodrigues vector norm) from its per-camera median across sessions. Dashed red lines mark a ± 5° stability threshold. Across all datasets, cameras exhibited minor session-to-session displacement and orientation drift, yet pose estimation accuracy remained consistent, demonstrating that our pipeline is robust to the rig perturbations typical of real experimental settings.

## Footnotes

1 https://github.com/tqxli/sdannce

2 Pretrained weights available at https://duke.app.box.com/s/2aw5r4hb3u57p1abt99n15f6hkl36×5k

3 https://github.com/tqxli/sdannce/blob/master/GUIDE.md

4 https://github.com/tqxli/sdannce/blob/master/demo/train_dannce.sh

